# Genome sequencing of *Musa acuminata* Dwarf Cavendish reveals a duplication of a large segment of chromosome 2

**DOI:** 10.1101/691923

**Authors:** Mareike Busche, Boas Pucker, Prisca Viehöver, Bernd Weisshaar, Ralf Stracke

**Affiliations:** Genetics and Genomics of Plants, Faculty of Biology & Center for Biotechnology (CeBiTec), Bielefeld University, Sequenz 1, 33615 Bielefeld, Germany

**Keywords:** banana, pan-genomics, small sequence variants, crop genome assembly

## Abstract

Different *Musa* species, subspecies, and cultivars are currently investigated to reveal their genomic diversity. Here, we compare the genome sequence of one of the commercially most important cultivars, *Musa acuminata* Dwarf Cavendish, against the Pahang reference genome assembly. Numerous small sequence variants were detected and the ploidy of the cultivar presented here was determined as triploid based on sequence variant frequencies. Illumina sequence data also revealed a duplication of a large segment on the long arm of chromosome 2 in the Dwarf Cavendish genome. Comparison against previously sequenced cultivars provided evidence that this duplication is unique to Dwarf Cavendish. Although no functional relevance of this duplication was identified, this example shows the potential of plants to tolerate such aneuploidies.

---

Bananas (*Musa*) are monocotyledonous perennial plants. The edible fruit (botanically a berry) belongs to the most popular fruits in the world. In 2016, about 5.5 million hectares of land were used for the production of more than 112 million tons of bananas (FAO 2019). The majority of bananas were grown in Africa, Latin America, and Asia where they offer employment opportunities and are important export commodities (FAO 2019). Furthermore, with an annual *per capita* consumption of more than 200 kg in Rwanda and more than 100 kg in Angola, bananas provide food security in developing countries (FAO 2019; Arias *et al*. 2003). While plantains or cooking bananas are commonly eaten as a staple food in Africa and Latin America, the softer and sweeter dessert bananas are popular in Europe and Northern America. Between 1998 and 2000, around 47 % of the world banana production and the majority of the dessert banana production relied on the Cavendish subgroup of cultivars (Arias *et al*. 2003). Therein the Dwarf Cavendish banana (“Dwarf” refering to the height of the pseudostem, not to the fruit size) is one of the commercially most important cultivars, along with Grand Naine (“Chiquita banana”).

Although Cavendish bananas are almost exclusively traded internationally, numerous varieties are used for local consumption in Africa and Southeast Asia. Bananas went through a long domestication process which started at least 7,000 years ago (Denham *et al*. 2003). The first step towards edible bananas was interspecific hybridisation between subspecies from different regions, which caused incorrect meiosis and diploid gametes (Perrier *et al*. 2011). The diversity of edible triploid banana cultivars resulted from human selection and triploidization of *Musa acuminata* as well as *Musa balbisiana* (Perrier *et al*. 2011).

These exciting insights into the evolution of bananas were revealed by the analysis of genome sequences. Technological advances boosted sequencing capacities and allowed the (re-)sequencing of genomes from multiple subspecies and cultivars. *M. acuminata* can be divided into several subspecies and cultivars. The first *M. acuminata* (DH Pahang) genome sequence has been published in 2012 (D’Hont *et al*. 2012), many more genomes have been sequenced recently including: banksii, burmannica, zebrina (Rouard *et al*. 2018), malaccensis (SRR8989632, SRR6996493), Baxijiao (SRR6996491, SRR6996491), Sucrier : Pisang_Mas (SRR6996492). Additionally, the genome sequences of other *Musa* species, *M. balbisiana* (Davey *et al*. 2013), *M. itinerans* (Wu *et al*. 2016), and *M. schizocarpa* (Belser *et al*. 2018), have already been published.

Here we report about our investigation of the genome of *M. acuminata* Dwarf Cavendish, one of the commercially most important cultivars. We identified an increased copy number of a segment of the long arm of chromosome 2, indicating that this region was duplicated in one haplophase.

## Materials and Methods

### Plant material and DNA extraction

*Musa acuminata* Dwarf Cavendish tissue culture seedlings were obtained from FUTURE EXOTICS/SolarTek (Düsseldorf, Germany) (Figure 1). Plants were grown under natural daylight at 21°C. Genomic DNA was isolated from leaves following the protocol of Dellaporta *et al*. (1983).

**Figure 1:**
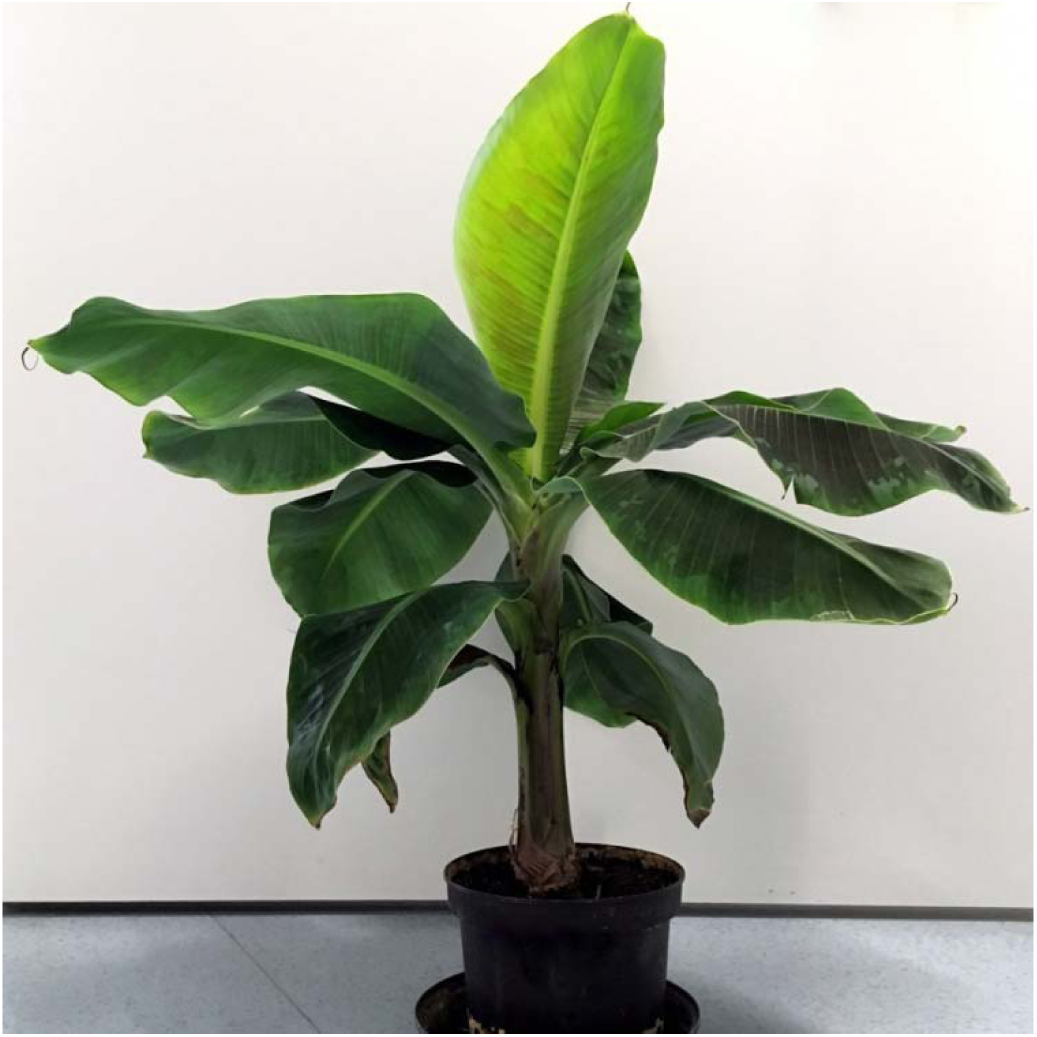
Nine month old *M. acuminata* Dwarf Cavendish plant.

### Library preparation and sequencing

Genomic DNA was subjected to sequencing library preparation via the TrueSeq v2 protocol as previously described (Pucker *et al*. 2016). Paired-end sequencing was performed on an Illumina HiSeq1500 and NextSeq500, respectively, resulting in 2×250 nt and 2×154 nt read data sets with an average Phred score of 38. These data sets provide 55x and 65.6x coverage, respectively, for the approximately 523 Mbp (D’Hont *et al*. 2012) haploid banana genome.

### Read mapping, variant calling, and variant annotation

All reads were mapped to the DH Pahang v2 reference genome sequence via BWA-MEM v0.7 (Li 2013) using –M to flag short hits for downstream filtering. This read mapping was analysed by the HaplotypeCaller of the Genome Analysis ToolKit (GATK) v3.8 (McKenna *et al*. 2010; Van der Auwera *et al*. 2013) to identify sequence variations in single nucleotides, called “single nucleotide variants” (SNVs), and also insertions/deletions (InDels). SNVs and InDels were called using the following filter rules in accordance with the GATK developer recommendation: ‘QD<2.0’, ‘FS>60.0’, and ‘MQ<40’ for SNVs and ‘QD<2.0’ and ‘FS>200.0’ for InDels. An InDel length cutoff of 100 bp was applied to restrict downstream analyses to a set of high quality variants called from 2×250nt reads. Only variants supported by at least five reads were kept. The resulting variant set was subjected to SnpEff (Cingolani *et al*. 2012) to assign predictions about the functional impact to the variants in the set. Variants with disruptive effects were selected using a customized Python script as described earlier (Pucker *et al*. 2016).

The genome-wide distribution of SNVs and InDels was assessed based on previously developed scripts (Baasner *et al*. 2019). The length distribution of InDels inside coding sequences was compared to the length distribution of InDels outside coding sequences using a customized Python script (Pucker *et al*. 2016).

### *De novo* genome assembly

Trimmomatic v0.38 (Bolger *et al*. 2014) was applied to remove low quality sequences (i.e. four consecutive bases below Phred 15) and remaining adapter sequences (based on similarity to all known Illumina adapter sequences). Different sets of trimmed reads were subjected to SOAPdenovo2 (Luo *et al*. 2012) for assembly using optimized parameters (Pucker *et al*. 2019) including avg_ins=600, asm_flags=3, rd_len_cutoff=300, pair_num_cutoff=3, and map_len=100. K-mer sizes ranged from 67 to 127 in steps of 10. Resulting assemblies were evaluated using previously described criteria (Pucker *et al*. 2019) including general assembly statistics (e.g. number of contigs, assembly size, N50, and N90) and a BUSCO (Benchmarking Universal Single-Copy Orthologs) v3 (Simão *et al*. 2015) assessment. Polishing was done by removing potential contaminations and adapters as described before (Pucker *et al*. 2019). The DH Pahang v2 assembly (D’Hont *et al*. 2012; Martin *et al*. 2016) was used in the contamination detection process to distinguish between *bona fide* banana contigs and sequences of unknown origin. Contigs with high sequence similarity to non-plant sequences were removed as previously described (Pucker *et al*. 2019). Remaining contigs were sorted based on the DH Pahang v2 reference genome sequence and concatenated to build pseudochromosomes to facilitate downstream analyses. A *de novo* Dwarf Cavendish assembly generated with a K-mer size of K=127 was choosen to give statistics.

## Data Availability Statement

Sequencing datasets were submitted to the European Nucleotide Archive (ERR3412983, ERR3412984, ERR3413471, ERR3413472, ERR3413473, ERR3413474). Python scripts are freely available on github (https://github.com/bpucker/banana). SNVs and InDels detected between the *M. acuminata* cultivars DH Pahang and Dwarf Cavendish are available in VCF format at http://doi.org/10.4119/unibi/2937972. The Dwarf Cavendish genome assembly is available in FASTA format at http://doi.org/10.4119/unibi/2937697.

## Supplementary Material

**File S1**. Per chromosome read coverage distribution of Dwarf Cavendish reads.

**File S2**. Comparison of SNVs in the duplicated segment on the long arm of chromosome 2 to all other SNVs in the genome. The higher read coverage at variants indicates a duplication of this region.

**File S3**. List of public genomic banana sequence read samples used for comparison against Dwarf Cavendish based on the DH Pahang reference.

**File S4**. Coverage plots of public genomic banana sequence read samples (File S1) for comparison against Dwarf Cavendish based on the DH Pahang reference. Samples are: *Musa acuminata, Musa acuminata* AYP_BOSN_r1, *Musa acuminata* ssp. banksii, *Musa acuminata* ssp. burmannica, *Musa acuminata* Cavendish BaXiJiao, *Musa acuminata* Gros Michel, *Musa acuminata* ssp. malaccensis, *Musa acuminata* Sucrier (Pisang Mas), *Musa acuminata* Sucrier (Pisang Mas 1998-2307), *Musa acuminata* ssp. zebrina (blood banana), *Musa balbisiana* Pisang Klutuk Wulung, *Musa itinerans, Musa schizocarpa*.

**File S5**. List of selected high impact variants between Dwarf Cavendish and DH Pahang with resulting effects predicted by SnpEff.

## Results and Discussion

### Structural variants

Mapping of *M. acuminata* Dwarf Cavendish reads against the DH Pahang v2 reference sequence assembly revealed several copy number variations in different parts of the genome (Figure 2, File S1). The most remarkable difference between the Dwarf Cavendish and Pahang genome sequence is the amplification of an about 6.2 Mbp continuous region (length deduced from the reference genome) on the long arm of chromosome 2 (Figure 2, File S1, S2). An investigation of allele frequencies in the duplicated segment on chromosome 2 revealed that this duplication originates from a haplophase with high similarity to the reference sequence (Figure 3). Such a duplication was not observed in any of the other publicly available genomic sequencing data sets when compared against the DH Pahang v2 genome sequence (File S3, S4). Apparently, read mapping also indicates at least four large scale deletions in Dwarf Cavendish compared to Pahang v2 on chromosomes 2, 4, 5 and 7 (Figure 2). However, analysis of the underlying sequence revealed long stretches of ambiguous bases (Ns) at these positions in the Pahang assembly as the cause for these pseudo low coverage regions.

**Figure 2:**
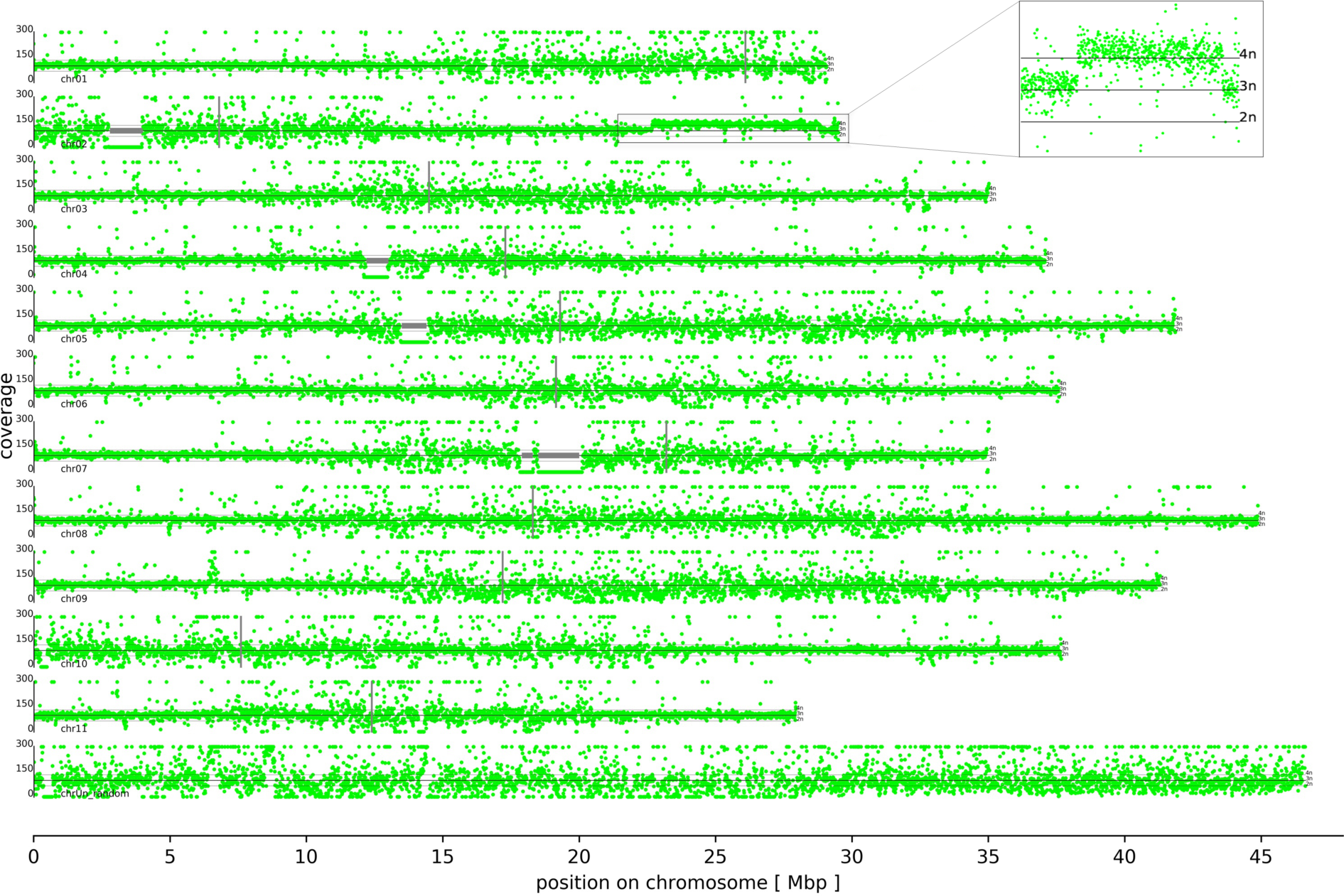
Coverage distribution. Chromosomes are ordered by increasing number with the north end on the left hand side. Centromere positions (D’Hont *et al*. 2012) are indicated by thin vertical grey lines. Mapping of *M. acuminata* Dwarf Cavendish reads against the DH Pahang v2 reference sequence assembly revealed a 6.2 Mbp tetraploid region on the long arm of chromosome 2 in Dwarf Cavendish (see enlarged box in the upper right). Apparent large scale deletions, indicated by regions with almost zero coverage, are technical artifacts caused by large stretches of ambiguous bases (Ns) in the Pahang assembly that cannot be covered by reads; these artifacts are marked with horizontal grey lines. Plots with higher per chromosome resolution data are presented in File S1.

### Ploidy of *M. acuminata* Dwarf Cavendish

Based on the coverage of small sequence variants (SNVs and InDels), the ploidy of Dwarf Cavendish was identified as triploid (Figure 3). Many heterozygous variant positions display a frequency of the reference allele close to 0.33 or close to 0.66. This fits the expectation for two copies of the reference allele and one copy of a different allele, or *vice versa*. Deviation from the precise values is explained by random fluctuation of the read distribution at the given position. Since the peak around 0.66 for the frequency of the allele identical to the reference is substantially higher than the peak around 0.33, it is reasonable to assume that two haplophases are very similar to the reference. The third haplophase is the one that contains more deviating positions and differs more from the reference. It is likely that reads of the divergent haplophase are mapped with a slightly reduced rate. This might explain why the peak at 0.66 is slightly more than twice the size of the peak at 0.33. In the duplicated segment on chromosome 2 the allele frequency peaks are shifted to 0.25 and 0.75 (Figure 3), indicating a tetraploid region with three haplophases identical to the reference and one haplophase divergent from the Pahang reference.

**Figure 3:**
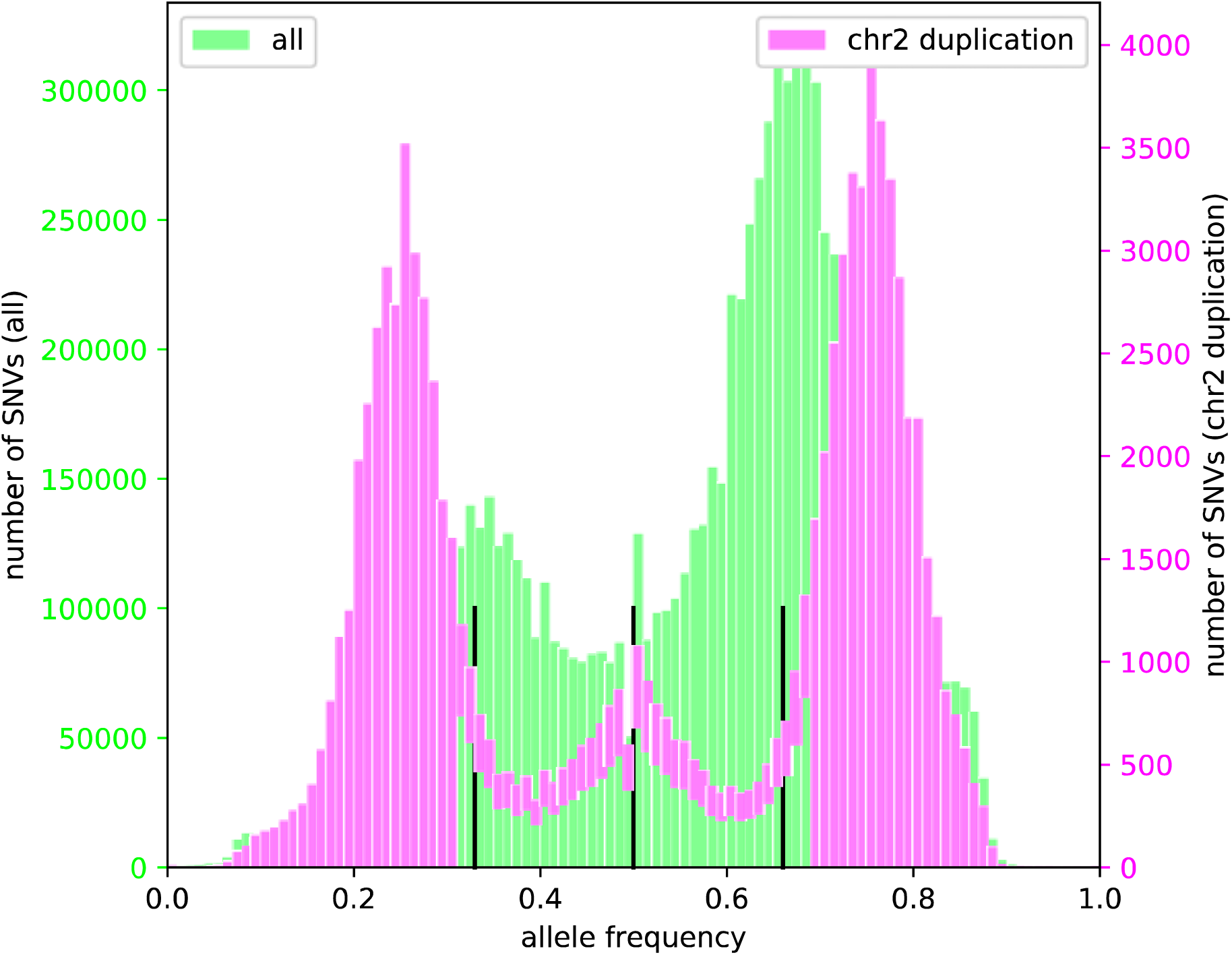
Allele frequency histogram. Visualisation of mapping results of Dwarf Cavendish Illumina reads against the Pahang v2 reference sequence, used for the determination of SNV frequencies. The frequencies of the reference allele at SNV positions are displayed here, excluding those positions at which the Pahang reference sequence deviates from an invariant sequence position of Dwarf Cavendish. Black vertical lines indicate allele frequencies of 0.33, 0.5, and 0.66, respectively. SNVs in the duplicated segment on the long arm of chromosome 2 (magenta) are distinguished from all other variants (lime). Within the duplicated segment on chromosome 2, the frequency of the reference alleles is often 0.75 or 0.25 indicating the presence of three similar alleles and one diverged allele.

To be able to test and prove or disprove hypotheses regarding differences of the haplophases of the Dwarf Cavendish genome, a high continuity phased assembly would be needed. Up-to-date long read sequencing technologies like Single Molecule Real-Time (Pacific Biosciences) or nanopore sequencing (Oxford Nanopore Technologies) in principle allow to generate such assemblies. However, successful phase separation currently requires tools like TrioCanu (Koren *et al*. 2018) which use Mendelian relationships between parents and F1 (i.e. crosses) for assignment of reads to phases. Generation of such datasets will be very difficult for banana and goes significantly beyond the scope of this study.

### Genome-wide distribution of small sequence variants

In total, 10,535,983 SNVs and 1,466,047 InDels were identified between the Dwarf Cavendish reads and the Pahang v2 assembly (see Data availability above). The genome-wide distribution of these variants is shown in Figure 4. As previously observed in other re-sequencing studies (Pucker *et al*. 2016), the number of SNVs exceeds the number of InDels substantially. Moreover, InDels are more frequent outside of annotated coding regions. Inside coding regions, InDels show an increased proportion of lengths which are divisible by 3, a bias introduced due to the avoidance of frameshifts.

**Figure 4:**
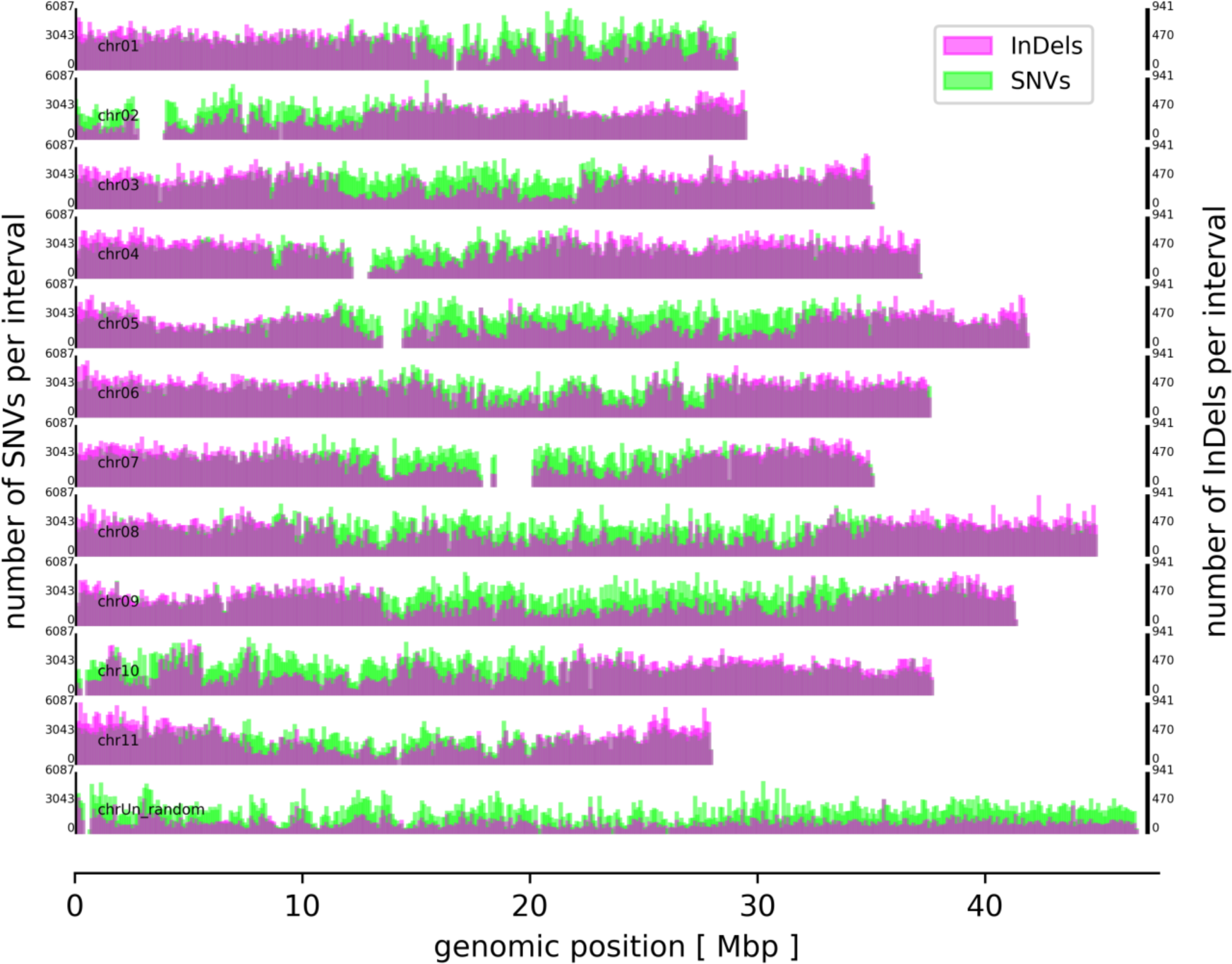
Genome-wide distribution of small sequence variants. SNVs (green) and InDels (magenta) distinguish Dwarf Cavendish from Pahang. Variants were counted in 100 kb windows and are displayed on two different y-axes to allow maximal resolution (Pucker *et al*. 2016).

SnpEff predicted 4,163 premature stop codons, 3,238 lost stop codons, and 8,065 frameshifts based on this variant set (File S5). Even given the larger genome size, these numbers are substantially higher than high impact variant numbers observed in re-sequencing studies of homozygous species before (Pucker *et al*. 2016; Xu *et al*. 2019). One explanation could be the presence of three alleles for each locus leading to compensation of disrupted alleles. Since banana plants are propagated vegetatively, breeders do not suffer inbreeding depressions.

### *De novo* genome assembly

To facilitate wet lab applications like oligonucleotide design and validation of amplicons, the genome sequence of Dwarf Cavendish was assembled *de novo*. The assembly comprises 256,523 scaffolds with an N50 of 5.4 kb (Table 1). Differences between the three haplophases are one possible explanation for the low assembly contiguity. The assembly size slightly exceeds the size of one haplotype. Due to the low contiguity of this assembly and only minimal above 50% complete BUSCOs (Benchmarking Universal Single-Copy Orthologs) (Simão *et al*. 2015), annotation was omitted. Nevertheless, we successfully used the produced genome assembly for primer design and detection of small sequence variants.

**Table 1:** *M. acuminata* Dwarf Cavendish *de novo* genome assembly statistics.

| Parameter | Value |
| --- | --- |
| Number of scaffolds | 256,523 |
| Maximal scaffold length | 240,314 bp |
| Assembly size | 963,409,601 bp (0.96 Gbp) |
| GC content | 38.78 % |
| N50 | 5,432 bp |
| N90 | 1,592 bp |

## Acknowledgments

We thank Joachim Weber for great technical assistance.

## Author contributions

BP, MB and RS planned the experiment. PV did the library preparation and sequencing. BP performed bioinformatic analyses. MB and BP wrote the initial draft. MB, BP, BW and RS revised the manuscript. All authors read and approved the final manuscript version.

## Notes

#### Summary of Updates

Updated link.

https://github.com/bpucker/banana

http://doi.org/10.4119/unibi/2937972

http://doi.org/10.4119/unibi/2937697

## Literature Cited

Arias, P., C. Dankers, P. Liu, and P. Pilkauskas, 2003 The World Banana Economy, 1985-2002. http://www.fao.org/3/y5102e/y5102e5100.htm.

Baasner, J.S., D. Howard, and B. Pucker, 2019 Influence of neighboring small sequence variants on functional impact prediction. bioRxiv doi: 10.1101/596718 (Preprint posted June 13, 2019).

Belser, C., B. Istace, E. Denis, M. Dubarry, F.C. Baurens et al., 2018 Chromosome-scale assemblies of plant genomes using nanopore long reads and optical maps. Natural Plants 4: 879–887.

Bolger, A.M., M. Lohse, and B. Usadel, 2014 Trimmomatic: a flexible trimmer for Illumina sequence data. Bioinformatics 30: 2114–2120.

Cingolani, P., A. Platts, L. Wang le, M. Coon, T. Nguyen et al., 2012 A program for annotating and predicting the effects of single nucleotide polymorphisms, SnpEff: SNPs in the genome of *Drosophila melanogaster* strain w1118; iso-2; iso-3. Fly (Austin) 6: 80–92.

D’Hont, A., F. Denoeud, J.M. Aury, F.C. Baurens, F. Carreel et al., 2012 The banana *(Musa acuminata)* genome and the evolution of monocotyledonous plants. Nature 488: 213–217.

Davey, M.W., R. Gudimella, J.A. Harikrishna, L.W. Sin, N. Khalid et al., 2013 A draft *Musa balbisiana* genome sequence for molecular genetics in polyploid, inter- and intra-specific Musa hybrids. BMC Genomics 14: 683.

Dellaporta, S.L., Wood, J. and Hicks, J.B., 1983 A plant DNA minipreparation: Version II. Plant Molecular Biology Reporter 1: 19–21.

Denham, T.P., S.G. Haberle, C. Lentfer, R. Fullagar, J. Field et al., 2003 Origins of agriculture at Kuk Swamp in the highlands of New Guinea. Science 301: 189–193.

FAO (Food and Agriculture Organization of the United Nations), 2019 FAOSTAT. http://www.fao.org/faostat/en/#data/QC.

Koren, S., A. Rhie, B.P. Walenz, A.T. Dilthey, D.M. Bickhart et al., 2018 *De novo* assembly of haplotype-resolved genomes with trio binning. Nature Biotechnology 36: 1174–1182.

Li, H., 2013 Aligning sequence reads, clone sequences and assembly contigs with BWA-MEM. 1303.3997v1302 (Preprint posted May 26, 2013).

Luo, R., B. Liu, Y. Xie, Z. Li, W. Huang et al., 2012 SOAPdenovo2: an empirically improved memory-efficient short-read de novo assembler. Gigascience 1: 18.

Martin, G., F.C. Baurens, G. Droc, M. Rouard, A. Cenci et al., 2016 Improvement of the banana *“Musa acuminata”* reference sequence using NGS data and semi-automated bioinformatics methods. BMC Genetics 17: 243.

McKenna, A., M. Hanna, E. Banks, A. Sivachenko, K. Cibulskis et al., 2010 The Genome Analysis Toolkit: a MapReduce framework for analyzing next-generation DNA sequencing data. Genome Research 20: 1297–1303.

Perrier, X., E. De Langhe, M. Donohue, C. Lentfer, L. Vrydaghs et al., 2011 Multidisciplinary perspectives on banana (Musa spp.) domestication. Proceedings of the National Academy of Sciences of the United States of America 108 (28):11311–11318.

Pucker, B., T. Feng, and S. Brockhington, 2019 Next generation sequencing to investigate genomic diversity in Caryophyllales. bioRxiv: doi: 10.1101/646133 (Preprint posted June 27, 2019).

Pucker, B., D. Holtgräwe, T. Rosleff Sörensen, R. Stracke, P. Viehöver et al., 2016 A *De novo* genome sequence assembly of the *Arabidopsis thaliana* accession Niederzenz-1 displays presence/absence variation and strong synteny. PLoS ONE 11: e0164321.

Rouard, M., G. Droc, G. Martin, J. Sardos, Y. Hueber et al., 2018 Three New genome assemblies support a rapid radiation in *Musa acuminata* (Wild Banana). Genome Biology and Evolution 10: 3129–3140.

Simão, F.A., R.M. Waterhouse, P. Ioannidis, E.V. Kriventseva, and E.M. Zdobnov, 2015 BUSCO: assessing genome assembly and annotation completeness with single-copy orthologs. Bioinformatics 31: 3210–3212.

Van der Auwera, G.A., M.O. Carneiro, C. Hartl, R. Poplin, G. Del Angel et al., 2013 From FastQ data to high confidence variant calls: the Genome Analysis Toolkit best practices pipeline. Current Protocols in Bioinformatics 11: 1110.

Wu, W., Y.L. Yang, W.M. He, M. Rouard, W.M. Li et al., 2016 Whole genome sequencing of a banana wild relative *Musa itinerans* provides insights into lineage-specific diversification of the Musa genus. Scientific Reports 6: 31586.

Xu, Y.C., X.M. Niu, X.X. Li, W. He, J.F. Chen et al., 2019 Adaptation and phenotypic diversification in Arabidopsis through loss-of-function mutations in protein-coding genes. The Plant Cell 31: 1012–1025.

