## Supplementary material for "Genome sequencing of *Musa acuminata* Dwarf Cavendish reveals a duplication of a large segment of chromosome 2": File S1

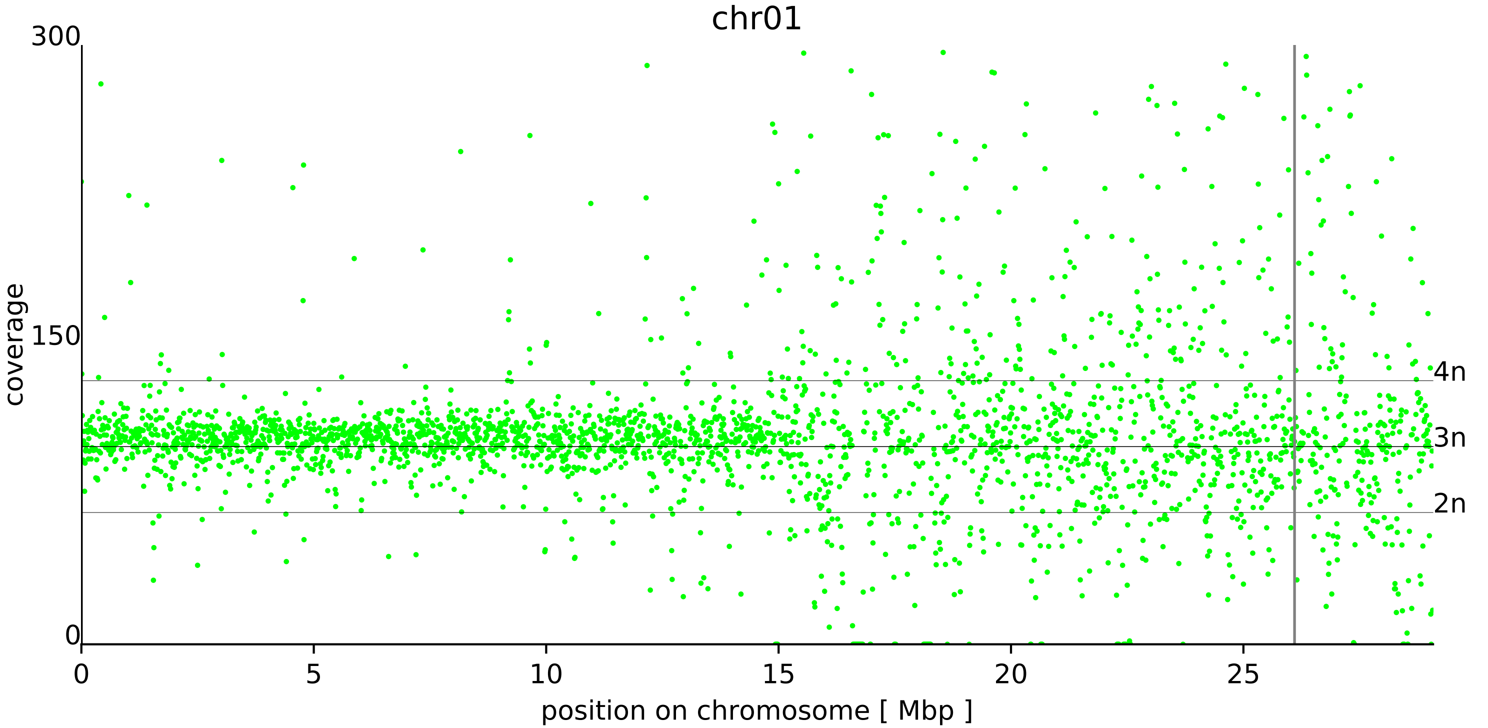
chr01
300
150
0
coverage
4n
3n
2n
0
5
10
15
20
25
position on chromosome [ Mbp ]

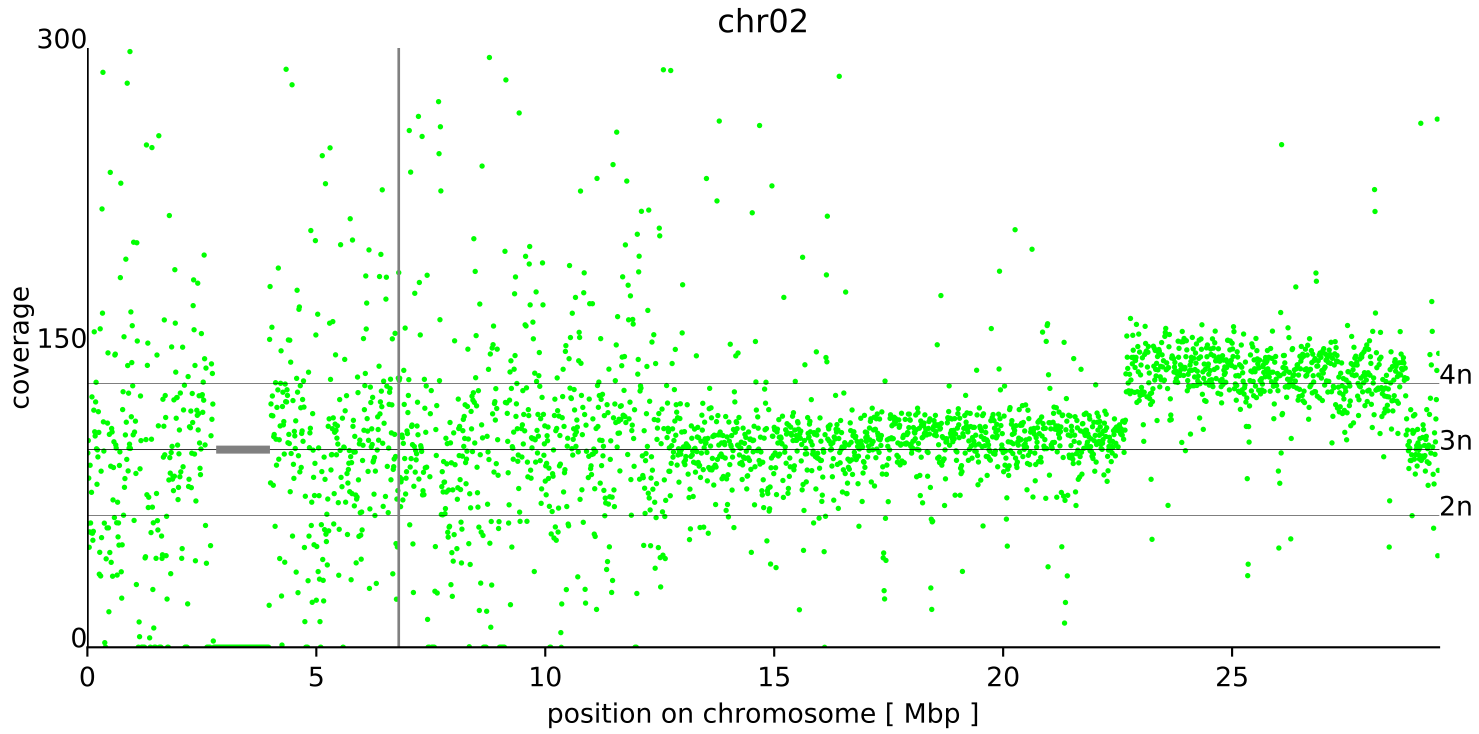
chr02
300
150
0
coverage
4n
3n
2n
0
5
10
15
20
25
position on chromosome [ Mbp ]

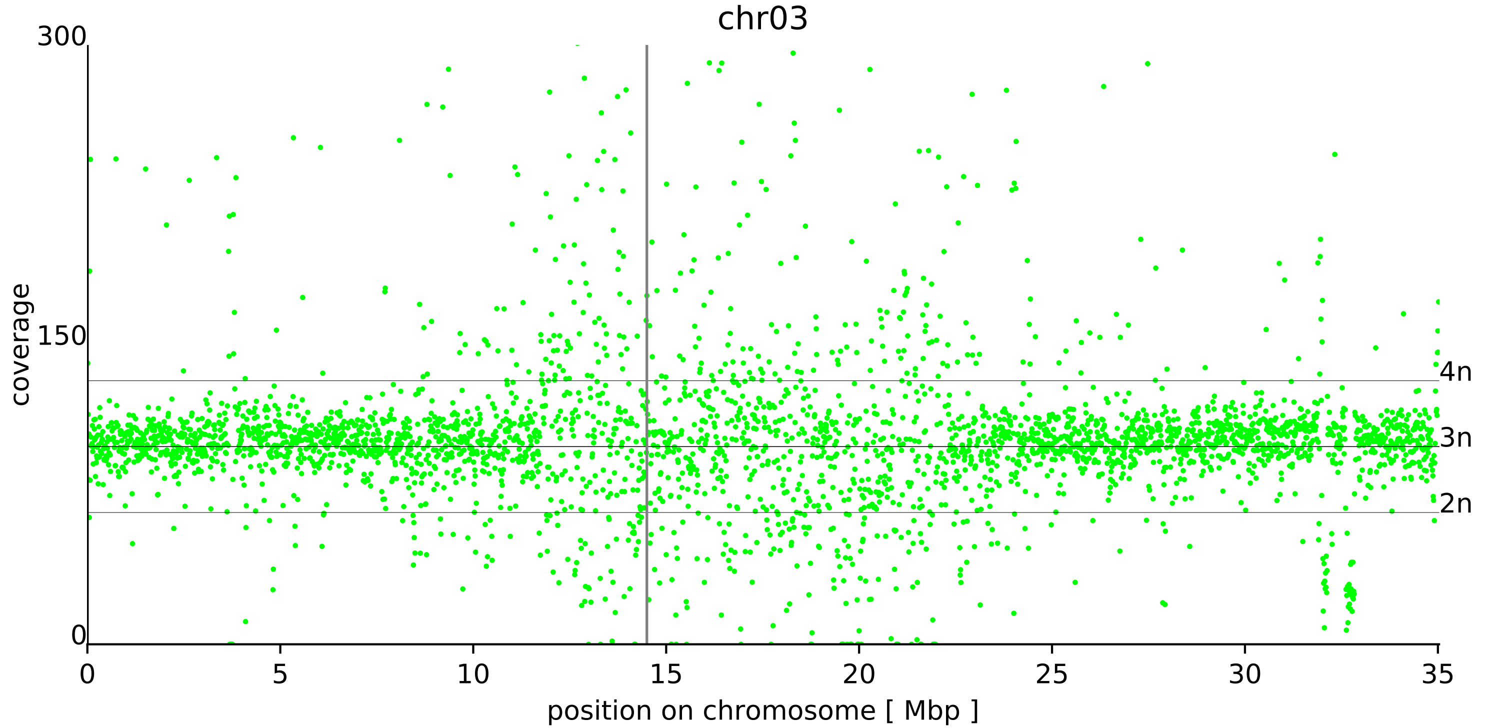

chr03
coverage
300
150
0
4n
3n
2n
0
5
10
15
20
25
30
35
position on chromosome [ Mbp ]

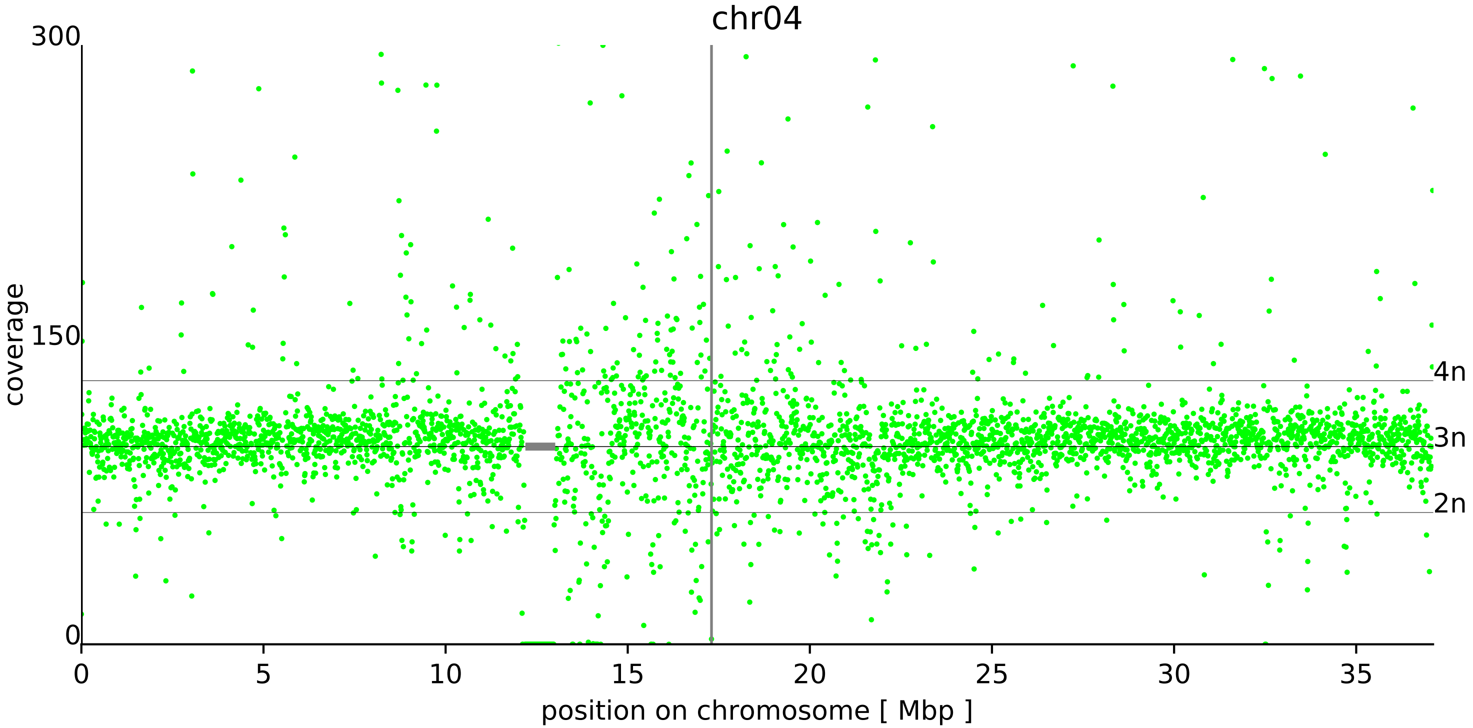

chr04
300
150
0
coverage
4n
3n
2n
0
5
10
15
20
25
30
35
position on chromosome [ Mbp ]

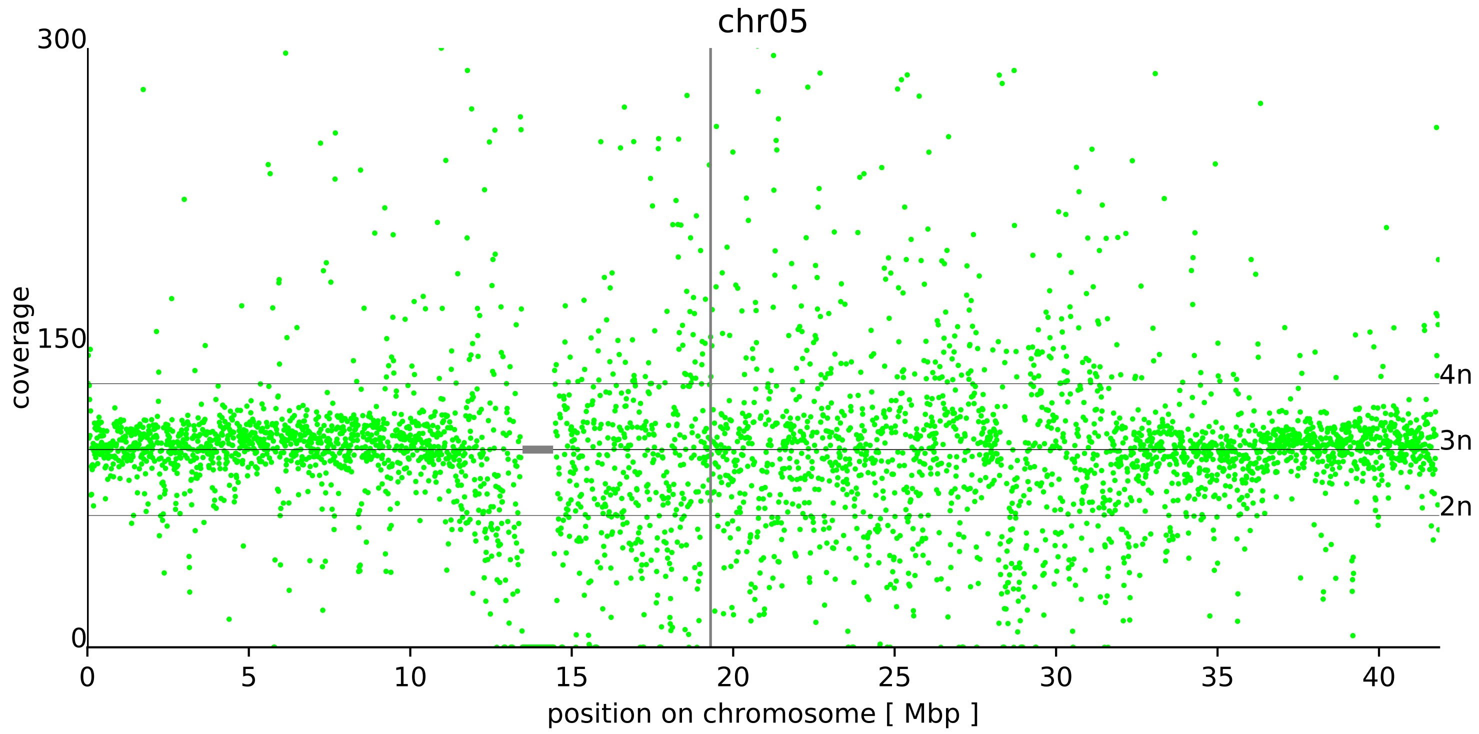
chr05
300
150
0
coverage
4n
3n
2n
0
5
10
15
20
25
30
35
40
position on chromosome [ Mbp ]

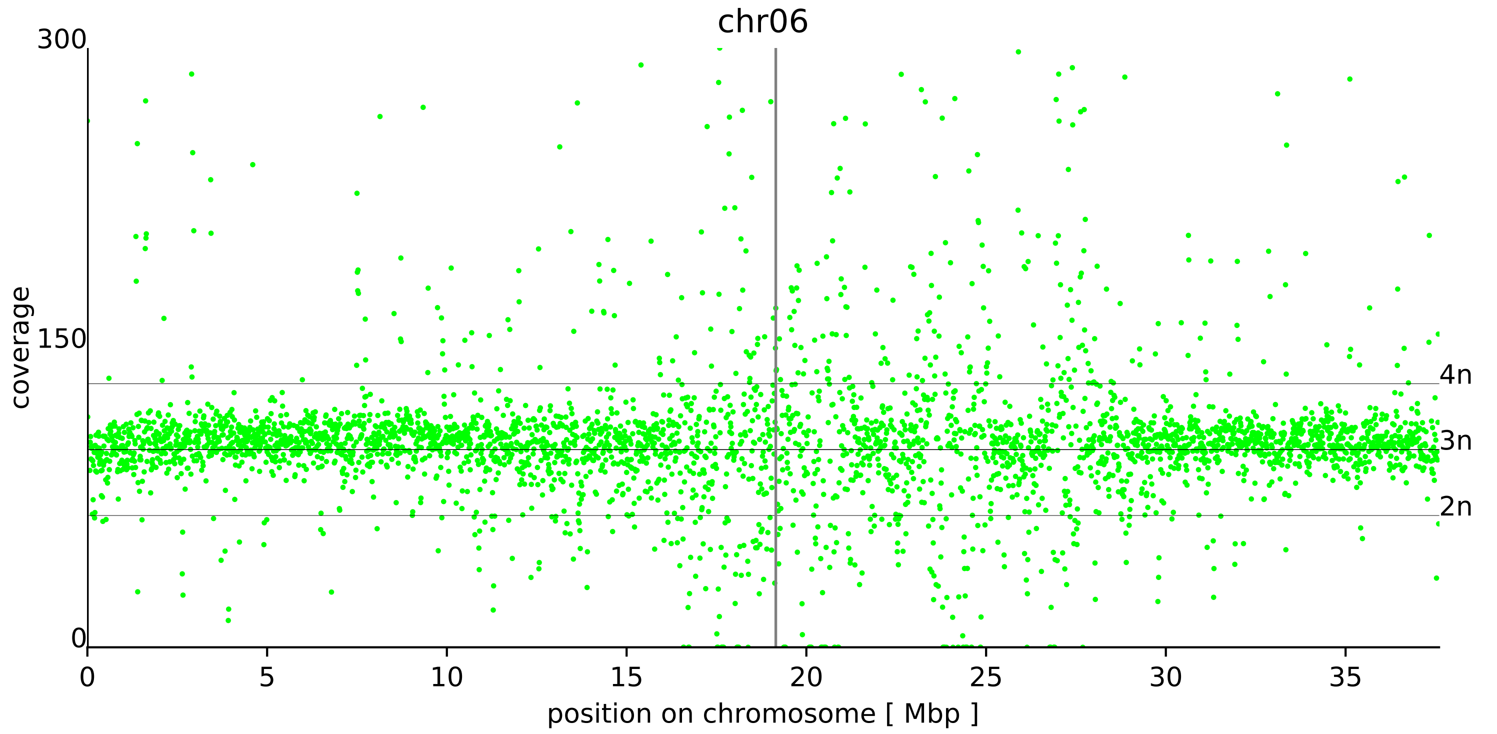
chr06
300
150
0
coverage
4n
3n
2n
0
5
10
15
20
25
30
35
position on chromosome [ Mbp ]

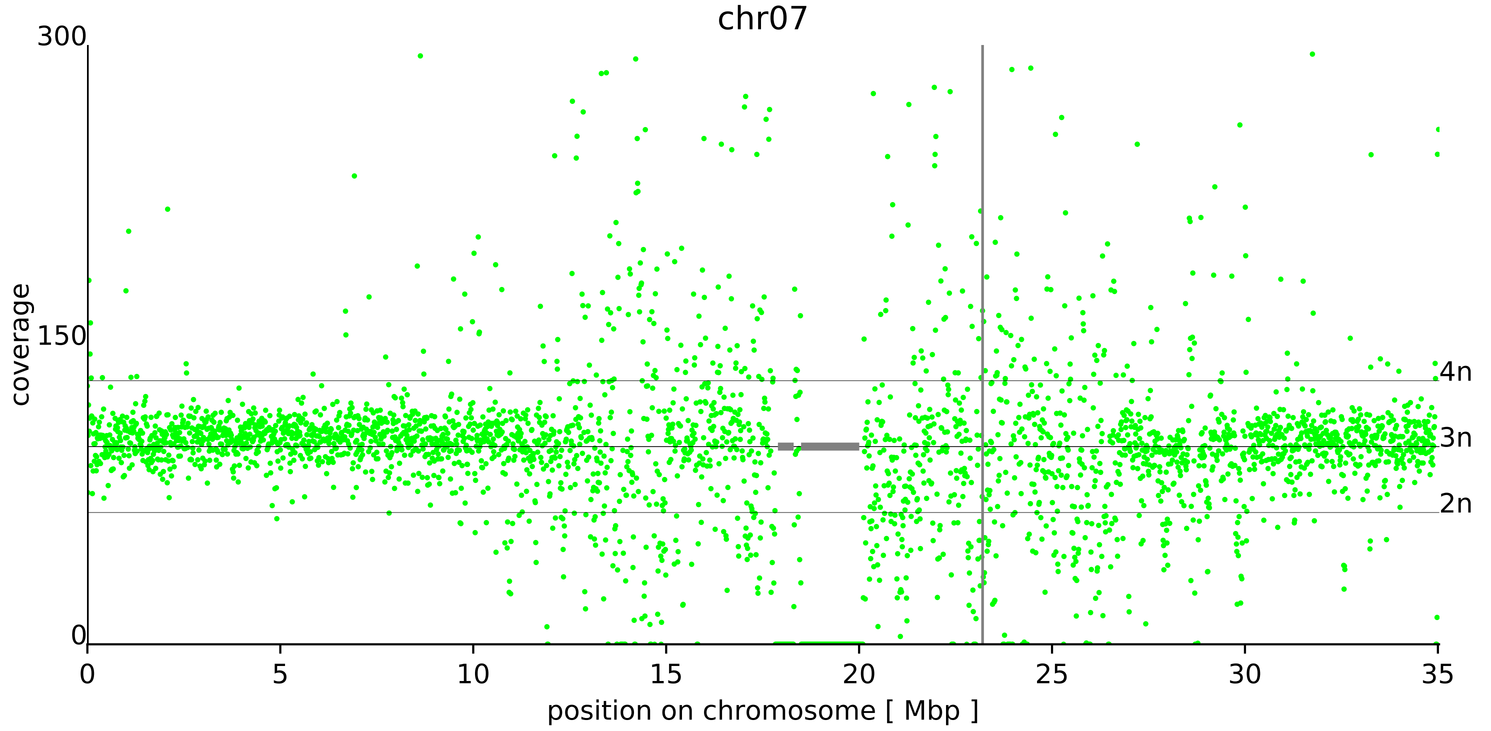

chr07
300
150
0
coverage
4n
3n
2n
0
5
10
15
20
25
30
35
position on chromosome [ Mbp ]

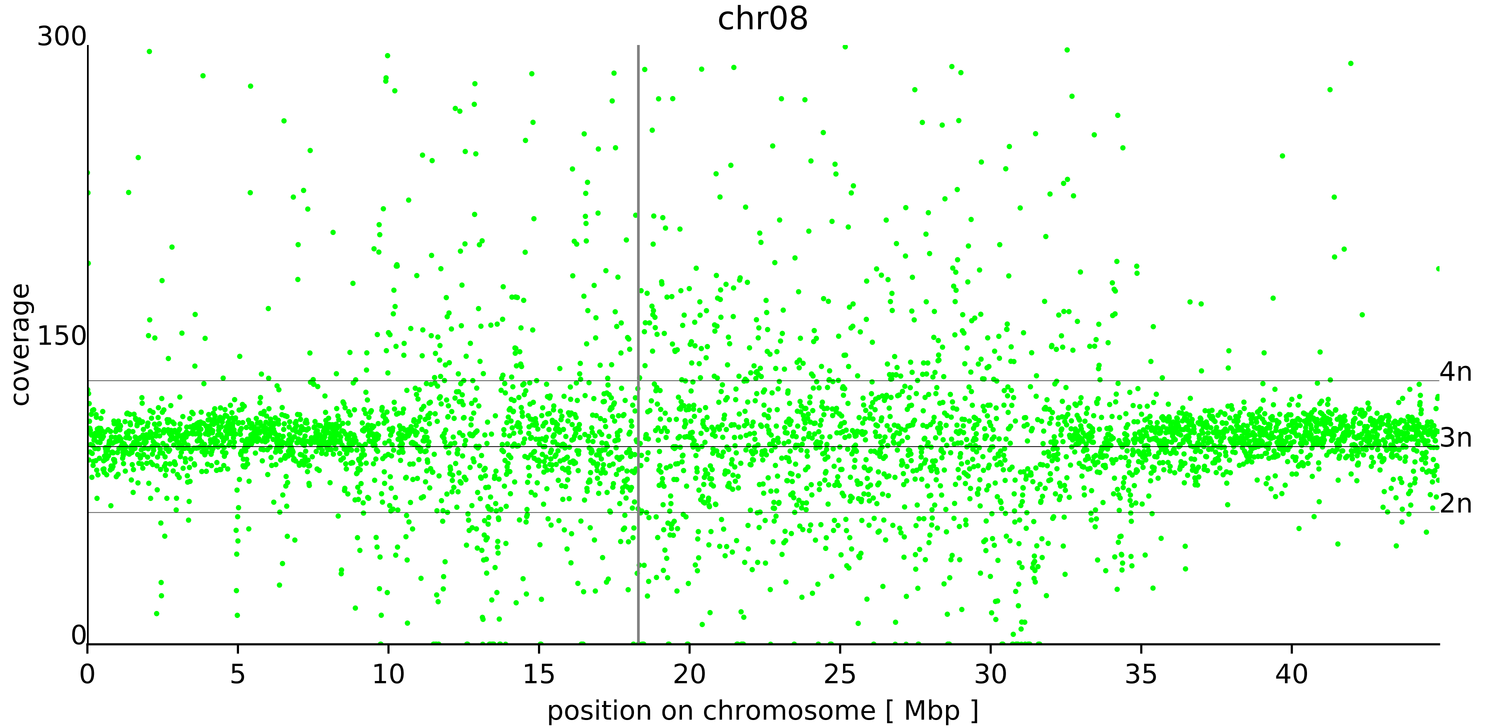

chr08
300
150
0
coverage
4n
3n
2n
0
5
10
15
20
25
30
35
40
position on chromosome [ Mbp ]

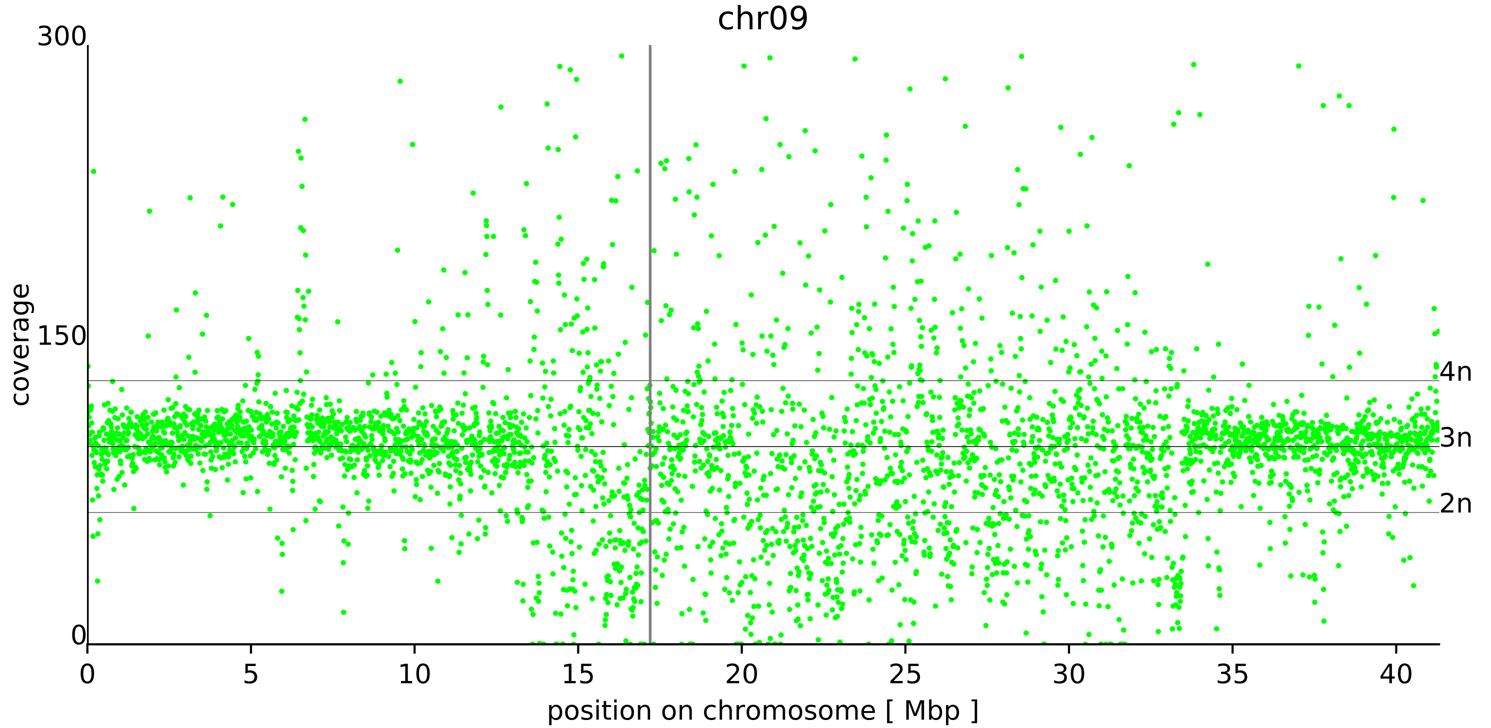

chr09
300
150
0
coverage
4n
3n
2n
0
5
10
15
20
25
30
35
40
position on chromosome [ Mbp ]

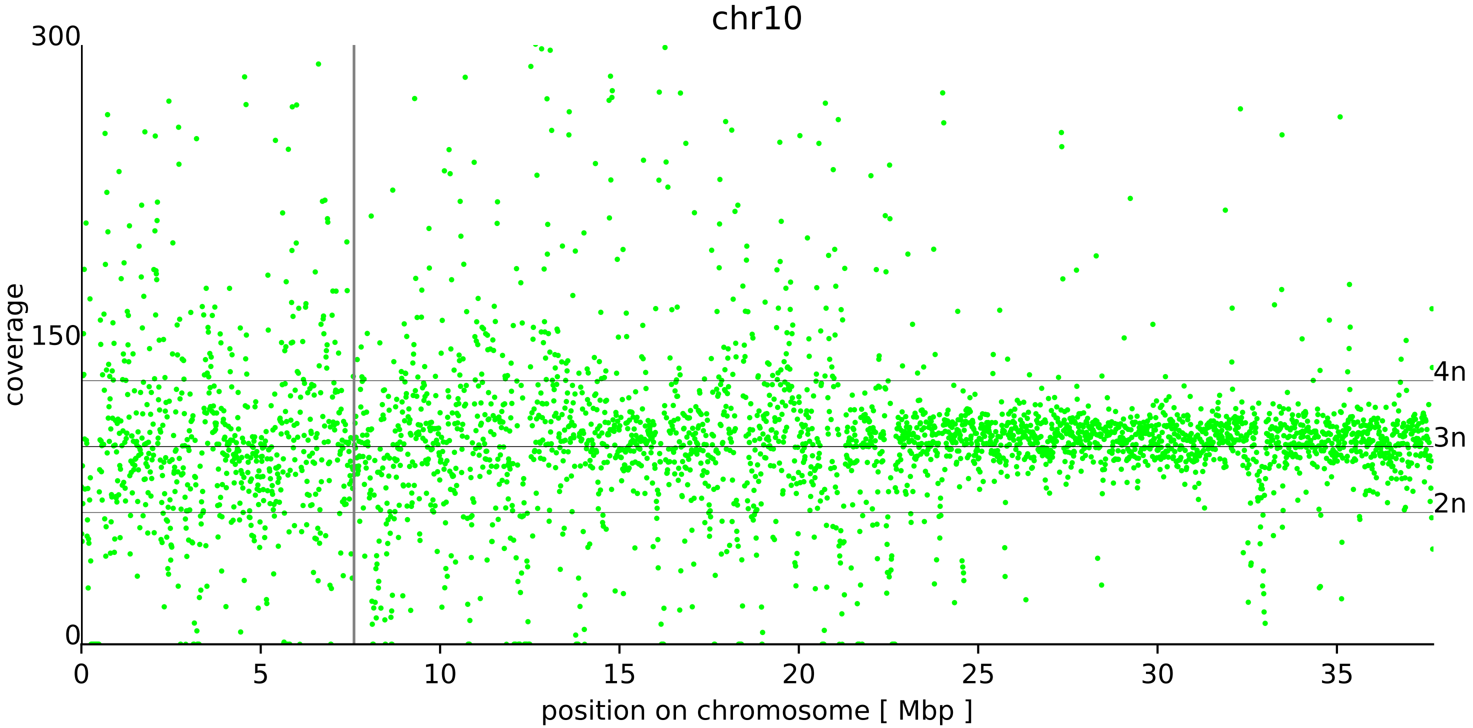

chr10
300
150
0
coverage
4n
3n
2n
0
5
10
15
20
25
30
35
position on chromosome [ Mbp ]

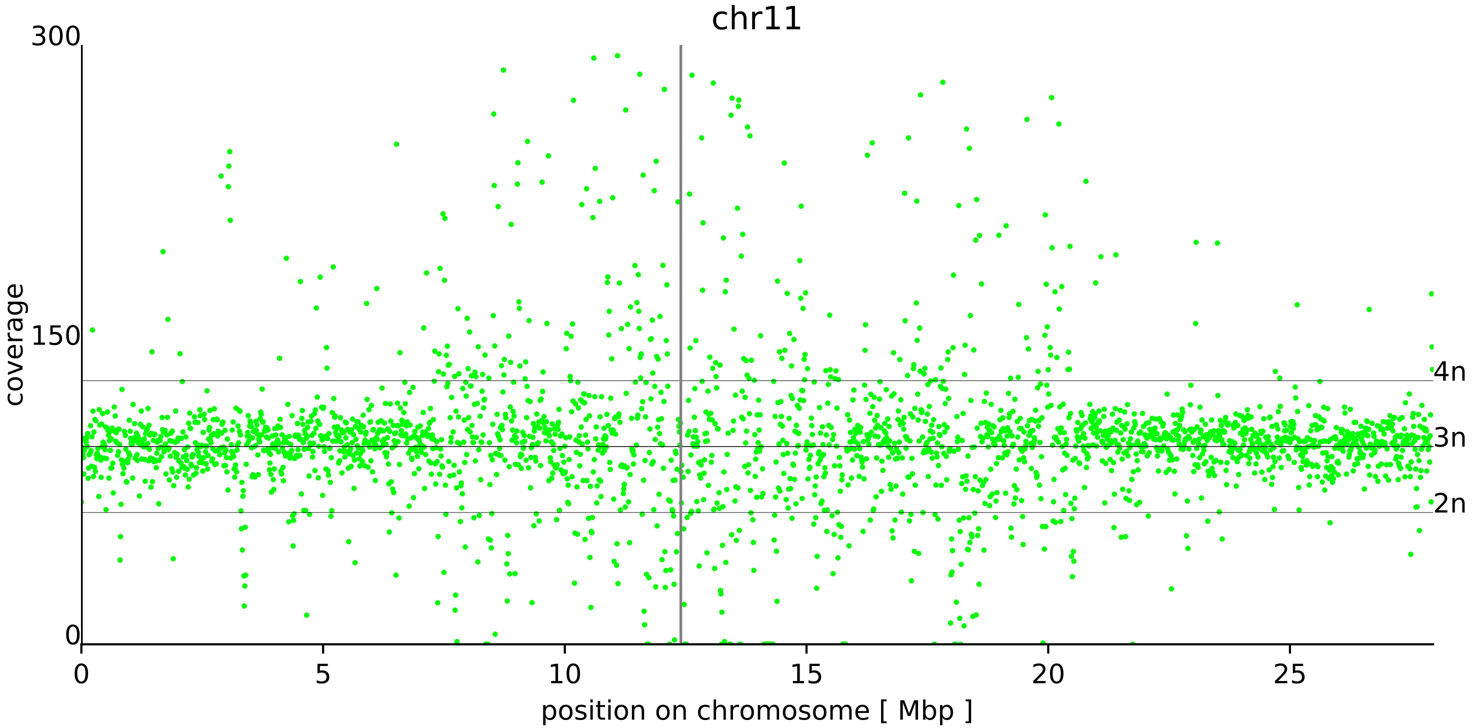
chr11
300
150
0
coverage
4n
3n
2n
0
5
10
15
20
25
position on chromosome [ Mbp ]

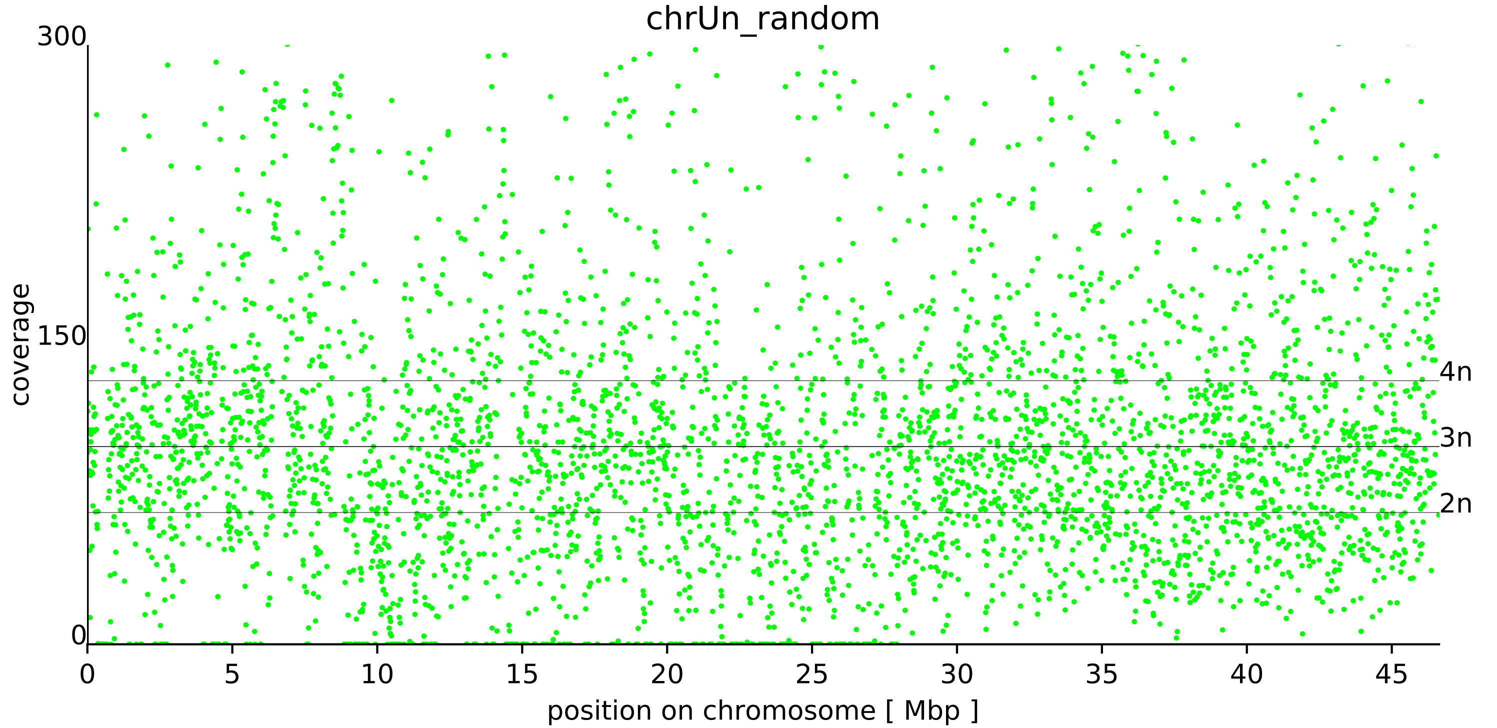

chrUn_random
300
150
0
coverage
4n
3n
2n
0
5
10
15
20
25
30
35
40
45
position on chromosome [ Mbp ]
