## Supplementary figures and images for "Genome sequencing of *Musa acuminata* Dwarf Cavendish reveals a duplication of a large segment of chromosome 2"

### File S2

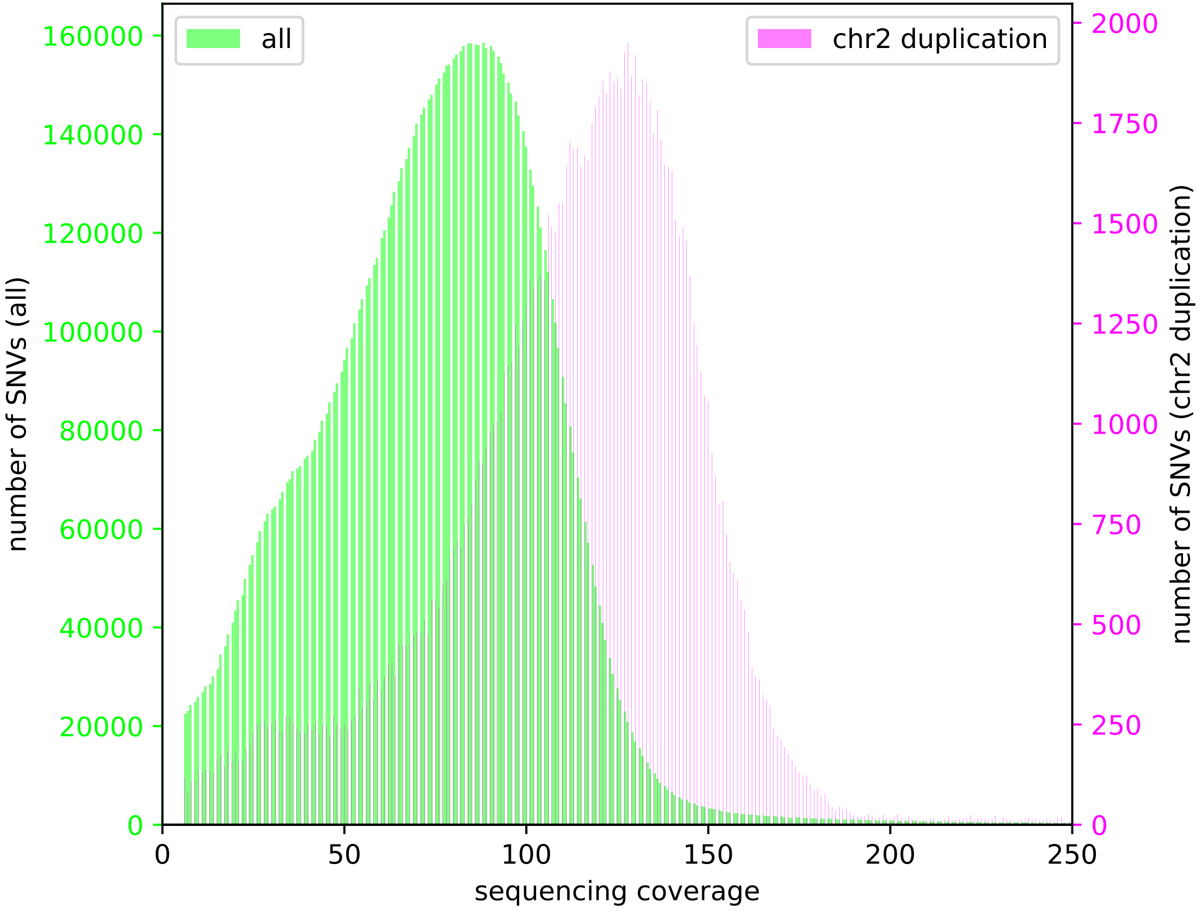
