## Supplementary material for "Genome sequencing of *Musa acuminata* Dwarf Cavendish reveals a duplication of a large segment of chromosome 2": File S4

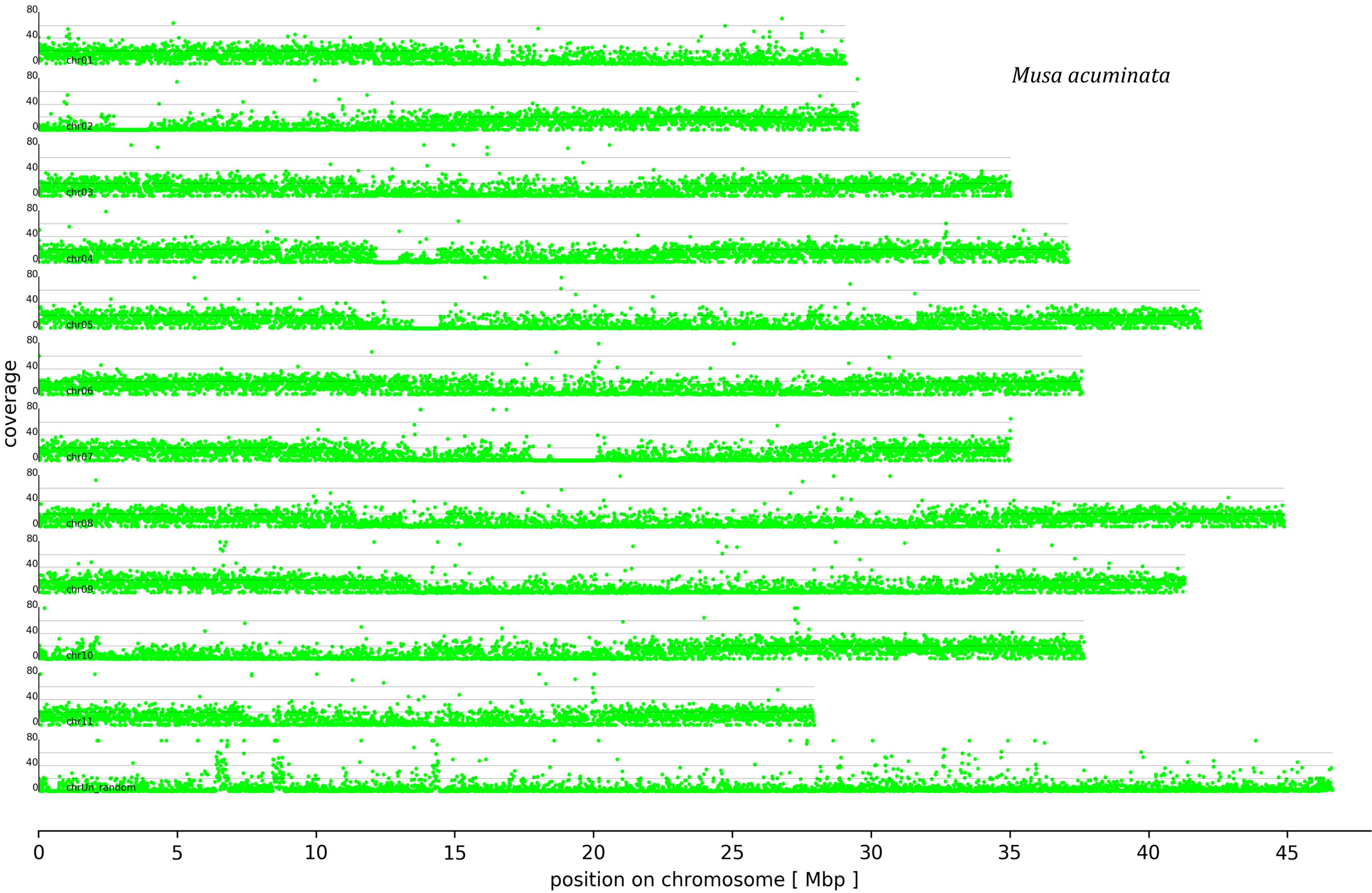
Musa acuminata
coverage
80
40
0
chr01
chr02
chr03
chr04
chr05
chr06
chr07
chr08
chr09
chr10
chr11
chrUn_random
0
5
10
15
20
25
30
35
40
45
position on chromosome [ Mbp ]

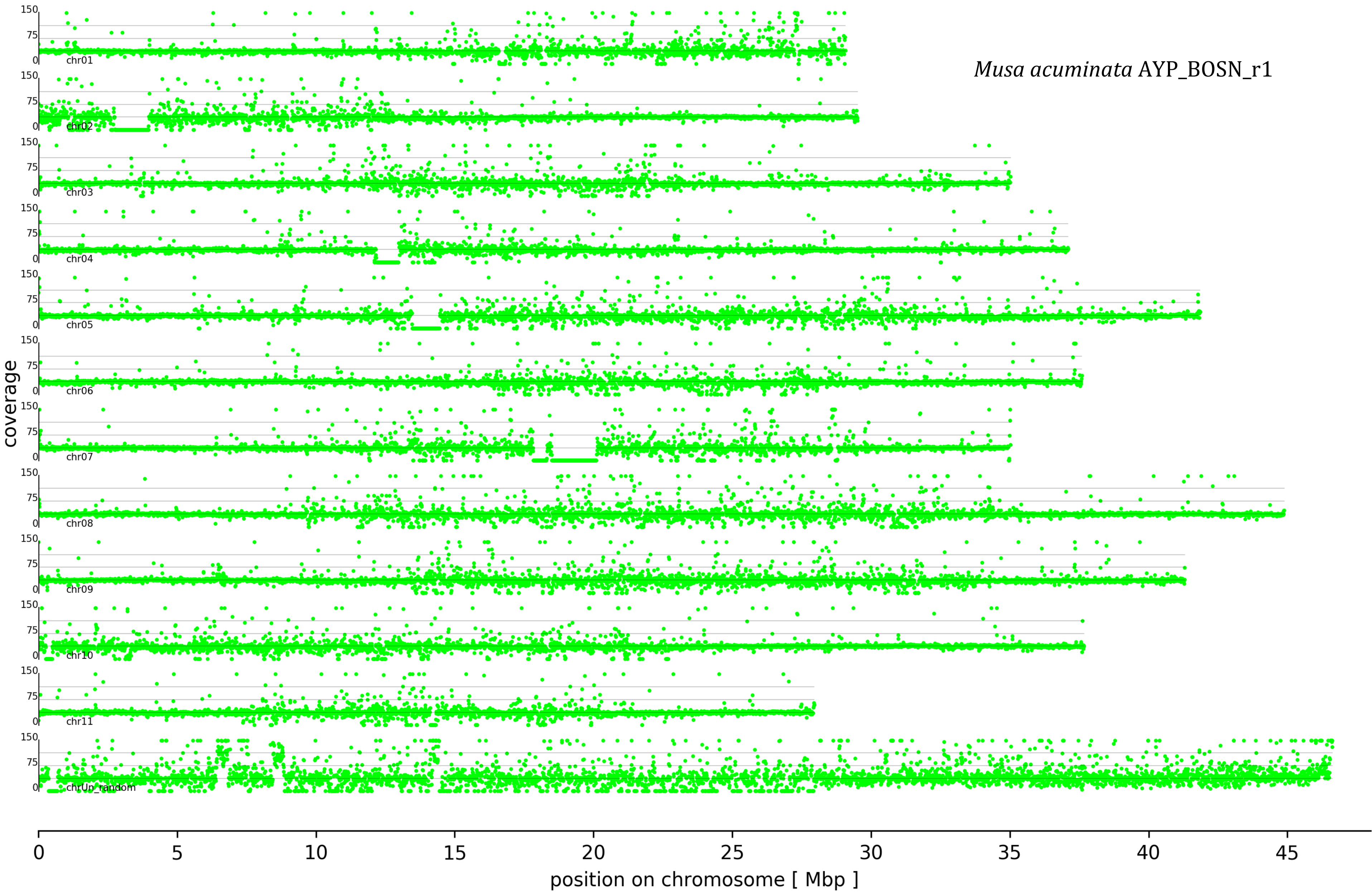
Musa acuminata AYP_BOSN_r1
coverage
150
75
0
chr01
chr02
chr03
chr04
chr05
chr06
chr07
chr08
chr09
chr10
chr11
chrUn_random
0
5
10
15
20
25
30
35
40
45
position on chromosome [ Mbp ]

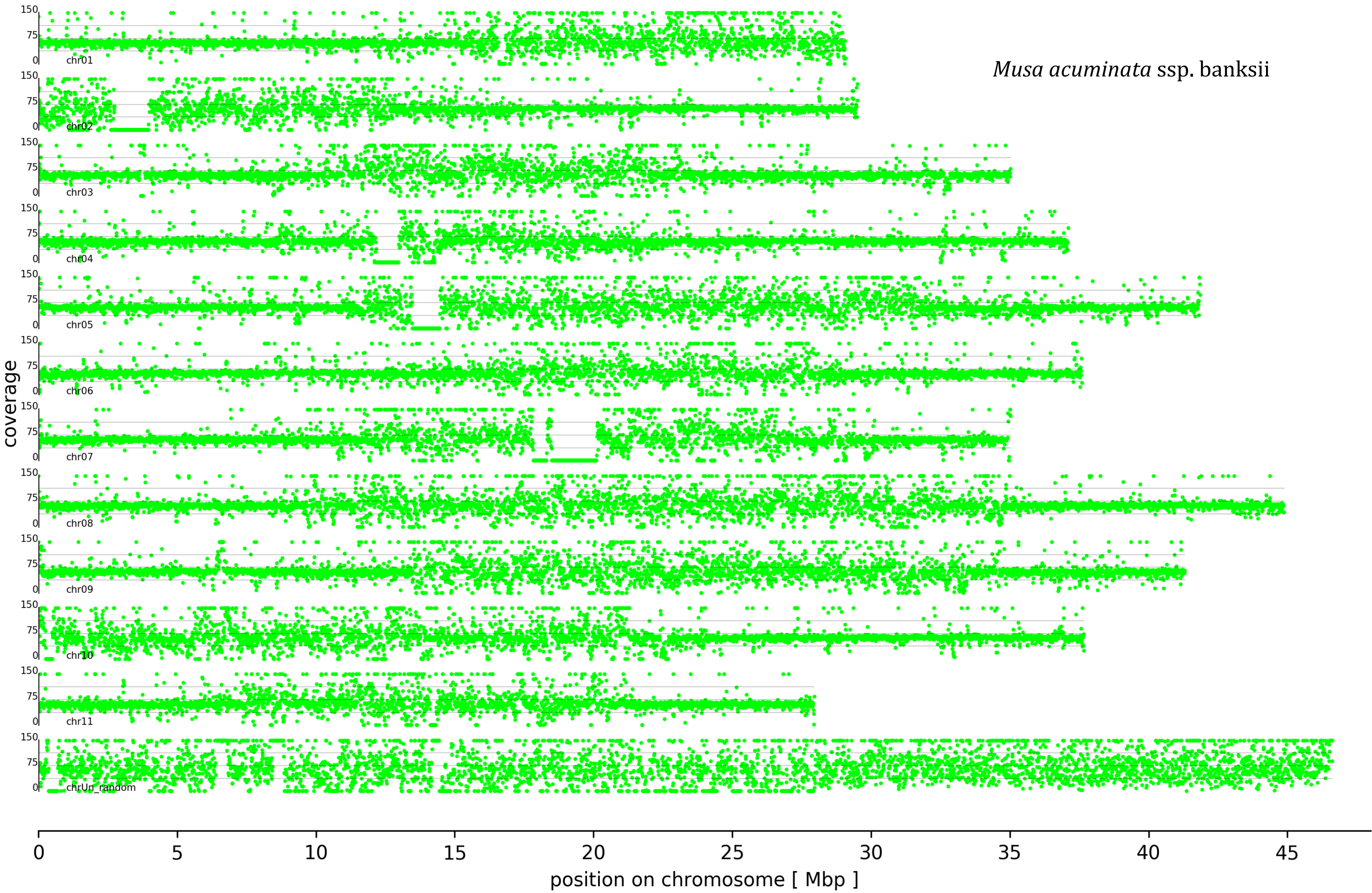

Musa acuminata ssp. banksii
coverage
150
75
0
chr01
chr02
chr03
chr04
chr05
chr06
chr07
chr08
chr09
chr10
chr11
chrUn_random
0
5
10
15
20
25
30
35
40
45
position on chromosome [ Mbp ]

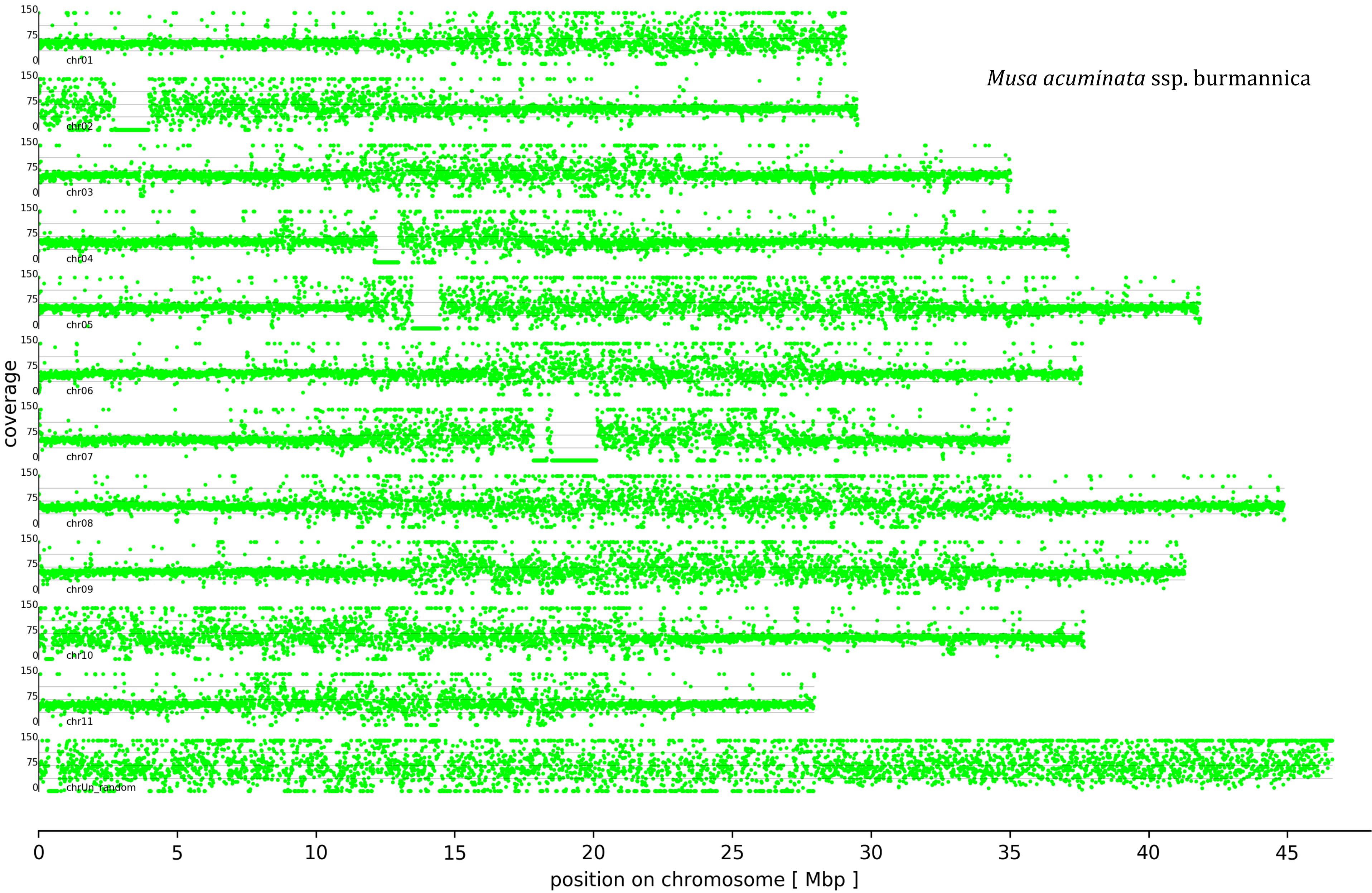
Musa acuminata ssp. burmannica
coverage
150
75
0
chr01
chr02
chr03
chr04
chr05
chr06
chr07
chr08
chr09
chr10
chr11
chrUn_random
0
5
10
15
20
25
30
35
40
45
position on chromosome [ Mbp ]

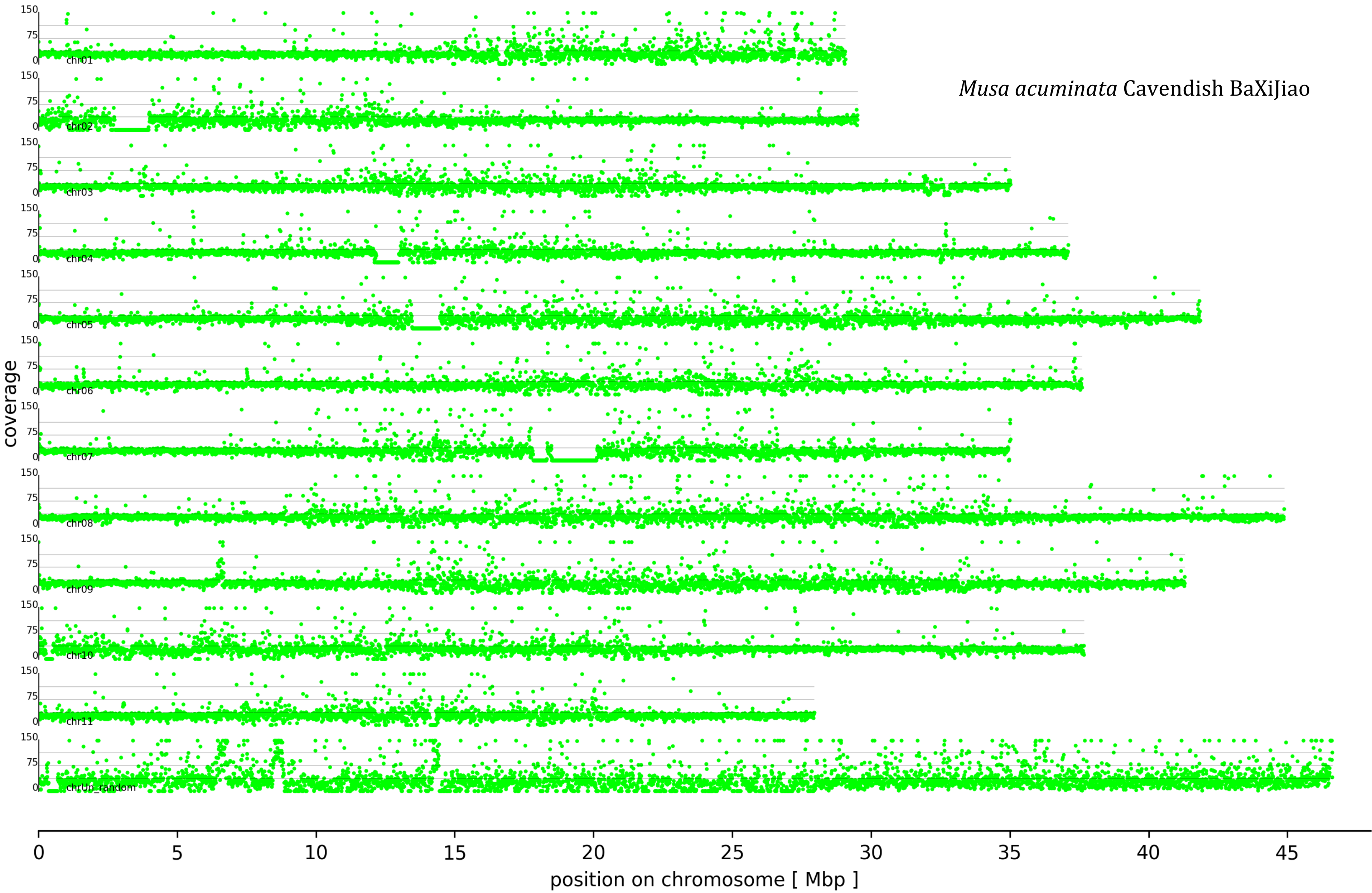

Musa acuminata Cavendish BaXiJiao
coverage
position on chromosome [ Mbp ]
chr01
chr02
chr03
chr04
chr05
chr06
chr07
chr08
chr09
chr10
chr11
chrUn_random
150
75
0
0
5
10
15
20
25
30
35
40
45

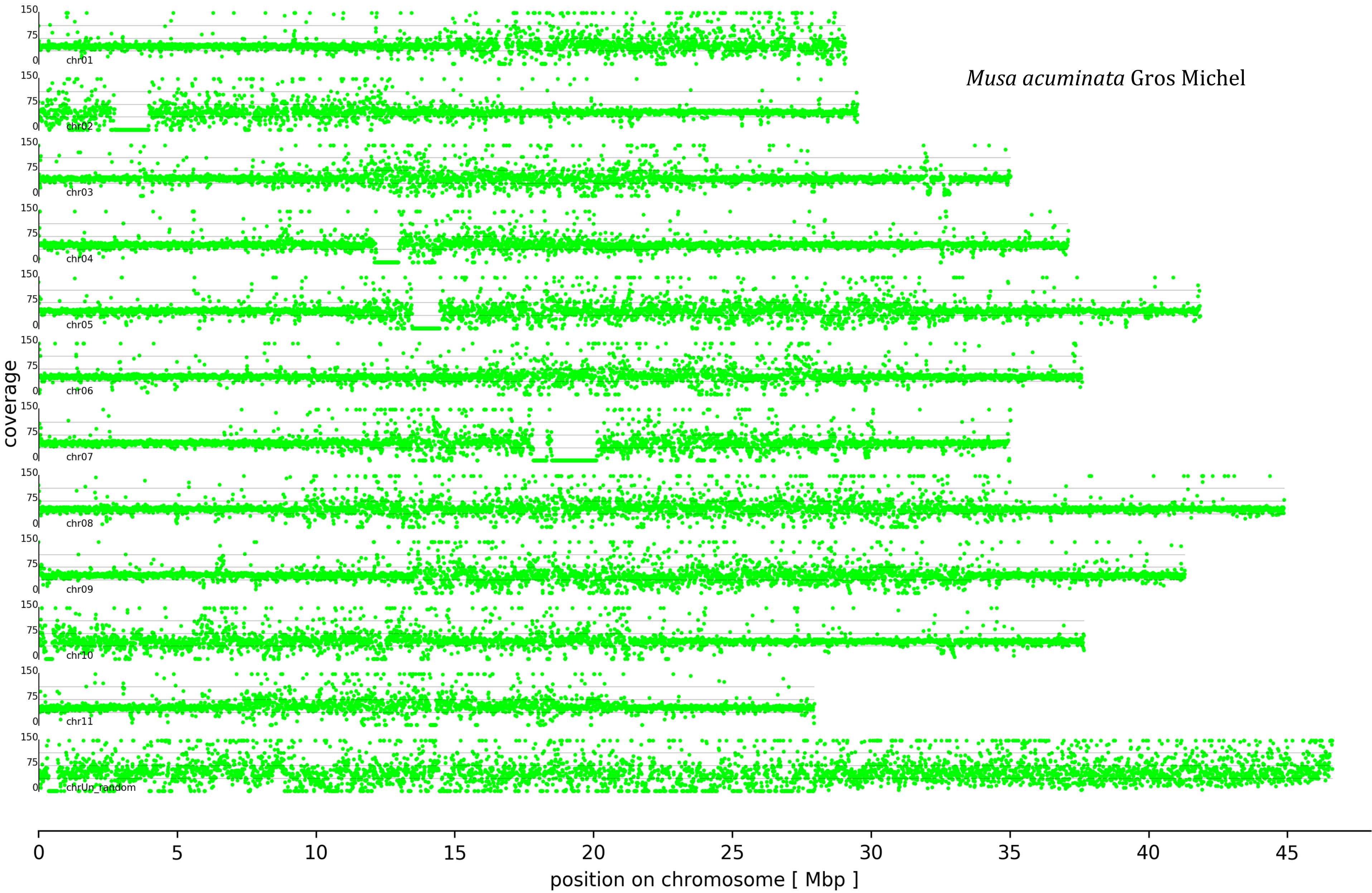
Musa acuminata Gros Michel
coverage
150
75
0
chr01
chr02
chr03
chr04
chr05
chr06
chr07
chr08
chr09
chr10
chr11
chrUn_random
0
5
10
15
20
25
30
35
40
45
position on chromosome [ Mbp ]

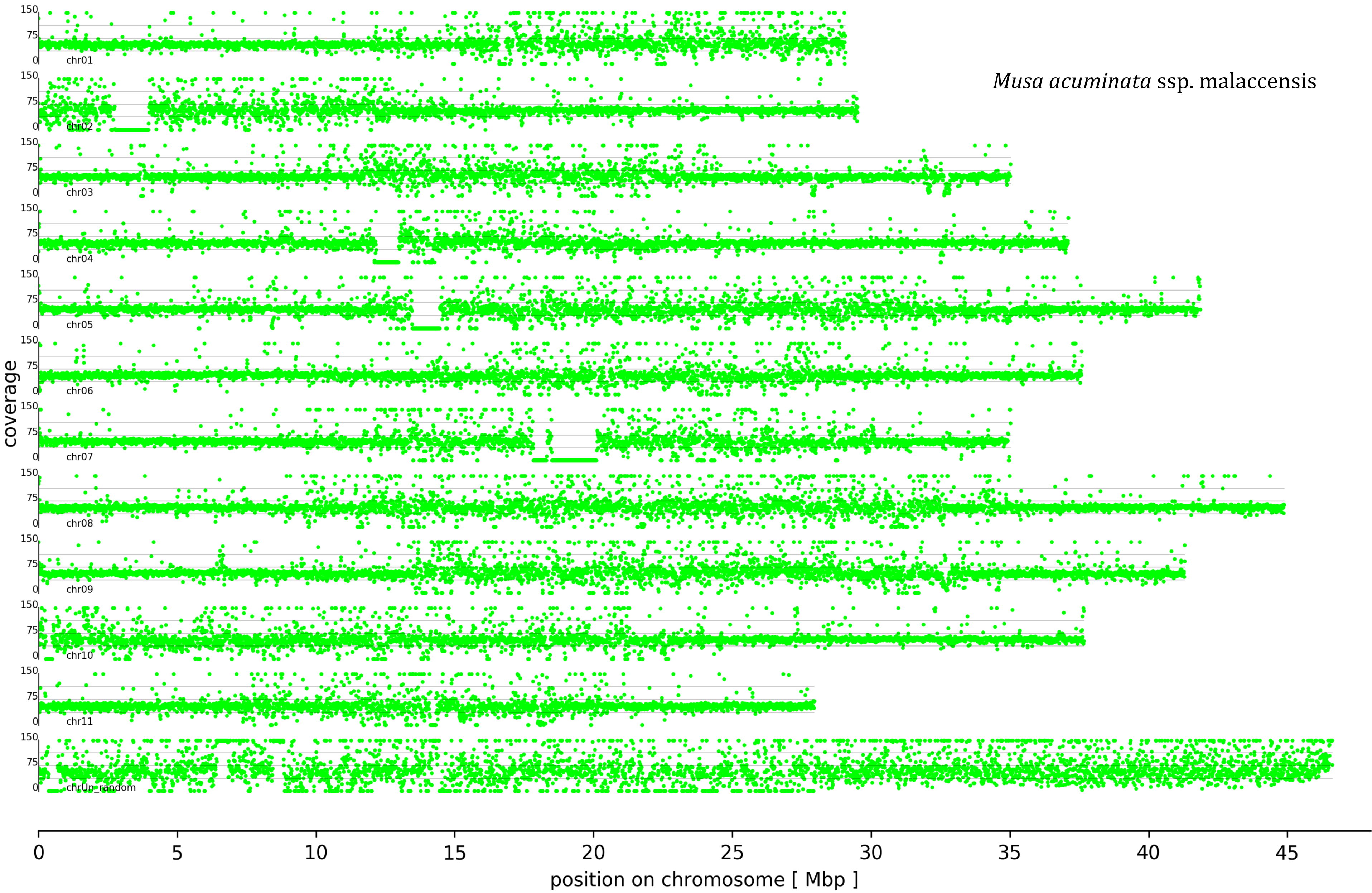

Musa acuminata ssp. malaccensis
coverage
150
75
0
chr01
chr02
chr03
chr04
chr05
chr06
chr07
chr08
chr09
chr10
chr11
chrUn_random
0
5
10
15
20
25
30
35
40
45
position on chromosome [ Mbp ]

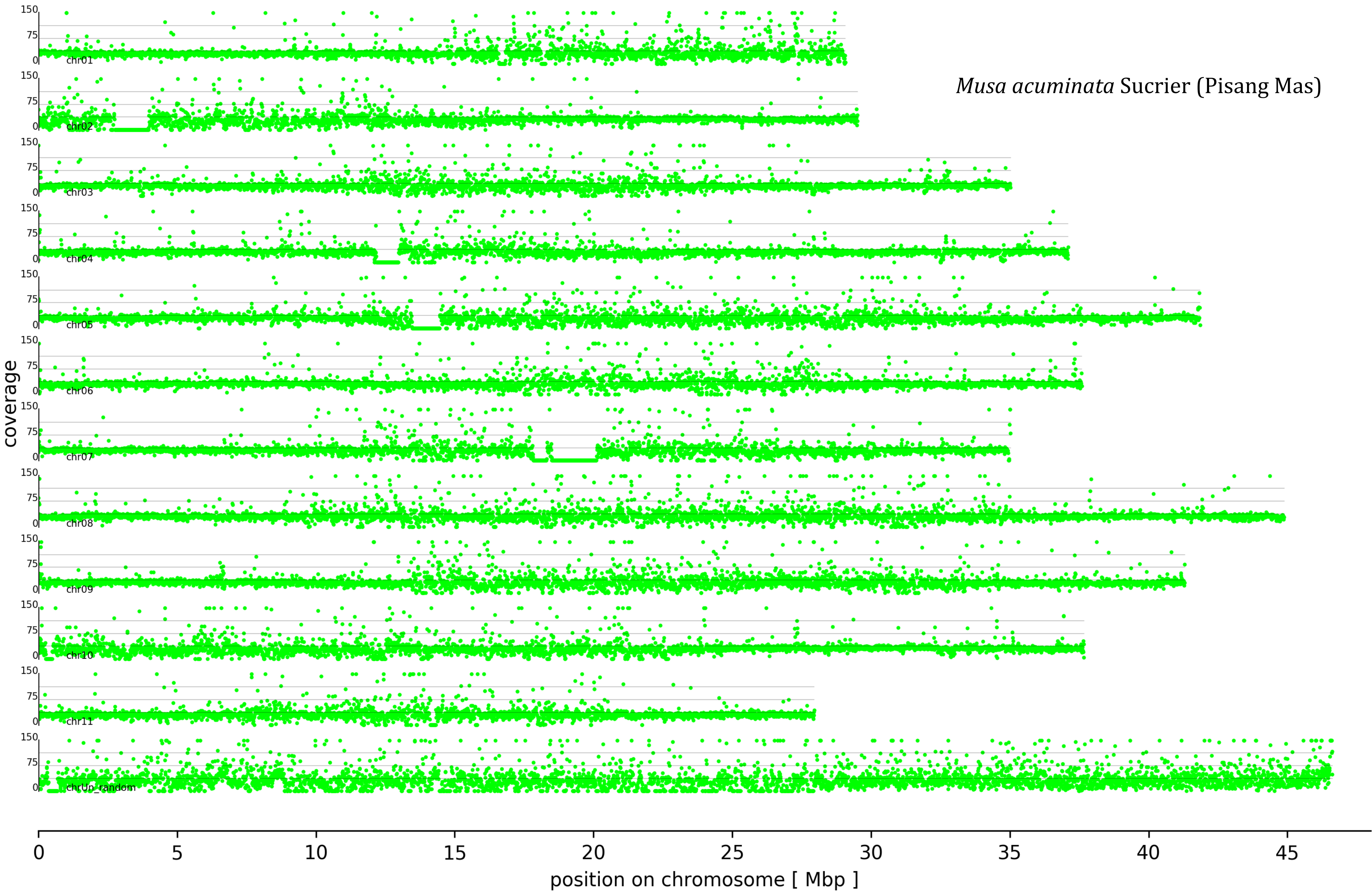

Musa acuminata Sucrier (Pisang Mas)
coverage
150
75
0
chr01
chr02
chr03
chr04
chr05
chr06
chr07
chr08
chr09
chr10
chr11
chrUn_random
0
5
10
15
20
25
30
35
40
45
position on chromosome [ Mbp ]

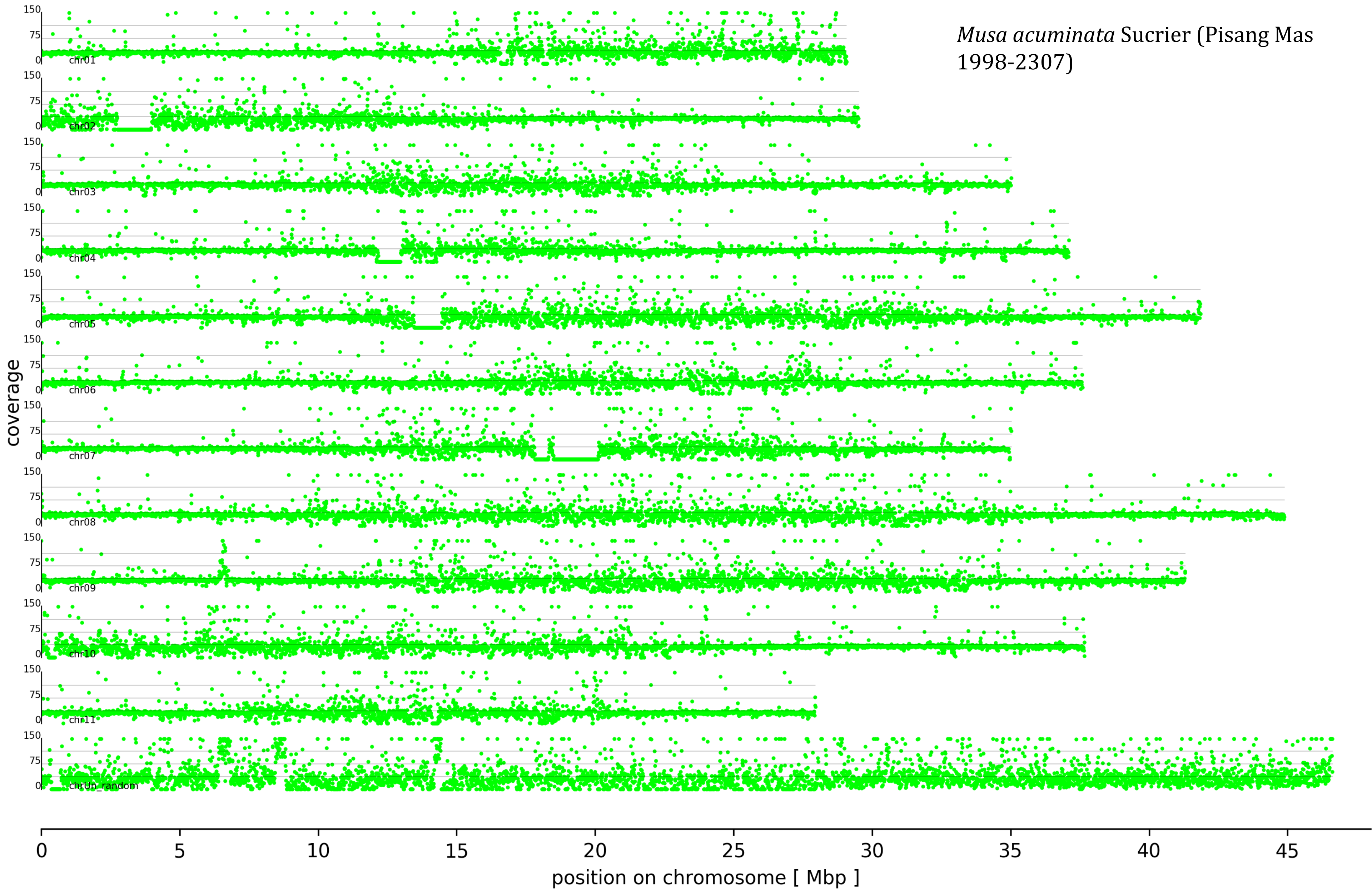

Musa acuminata Sucrier (Pisang Mas 1998-2307)
coverage
150
75
0
chr01
chr02
chr03
chr04
chr05
chr06
chr07
chr08
chr09
chr10
chr11
chrUn_random
0
5
10
15
20
25
30
35
40
45
position on chromosome [ Mbp ]

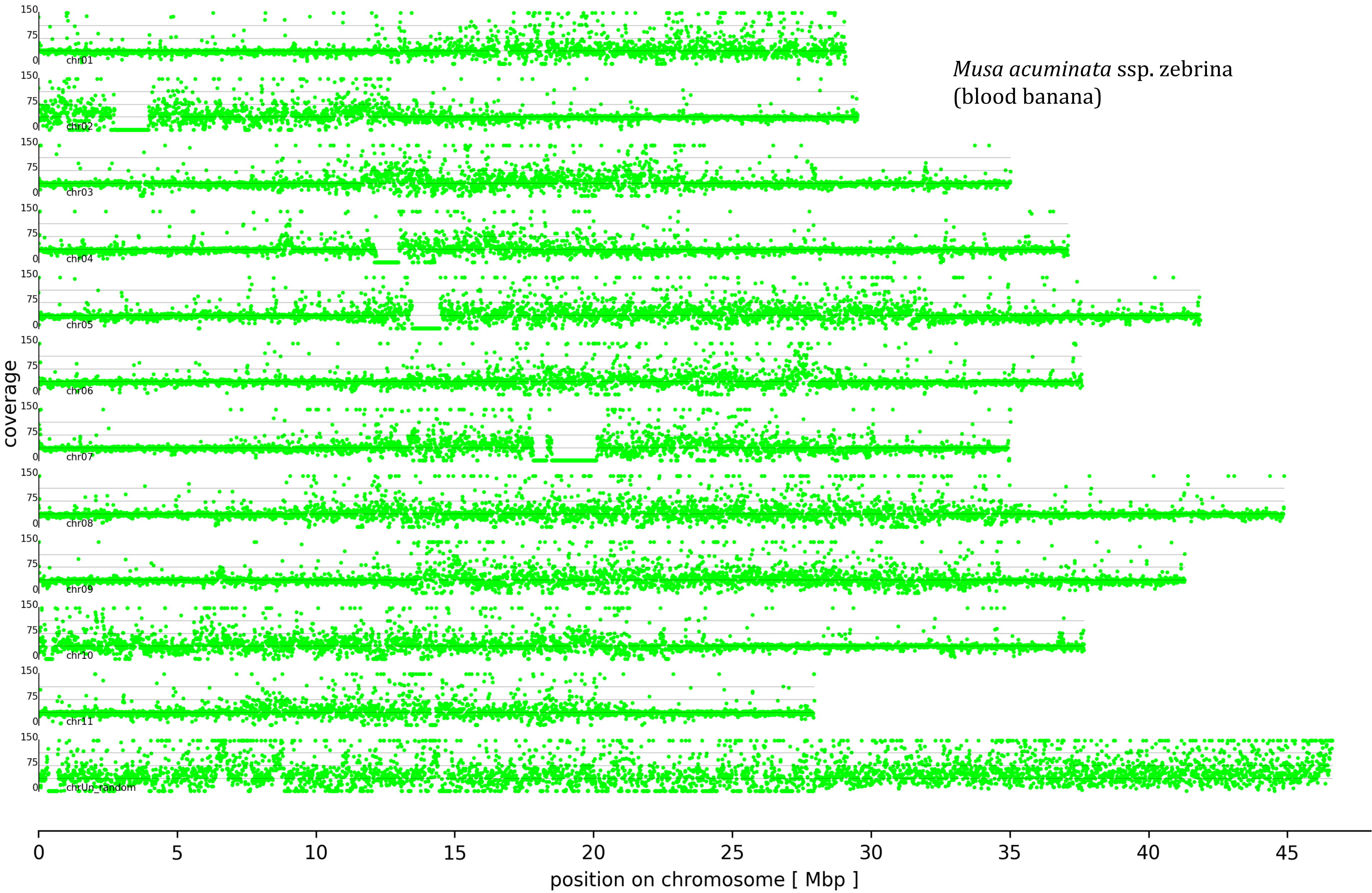

*Musa acuminata* ssp. zebrina
(blood banana)
coverage
150
75
0
chr01
chr02
chr03
chr04
chr05
chr06
chr07
chr08
chr09
chr10
chr11
chrUn_random
0
5
10
15
20
25
30
35
40
45
position on chromosome [ Mbp ]

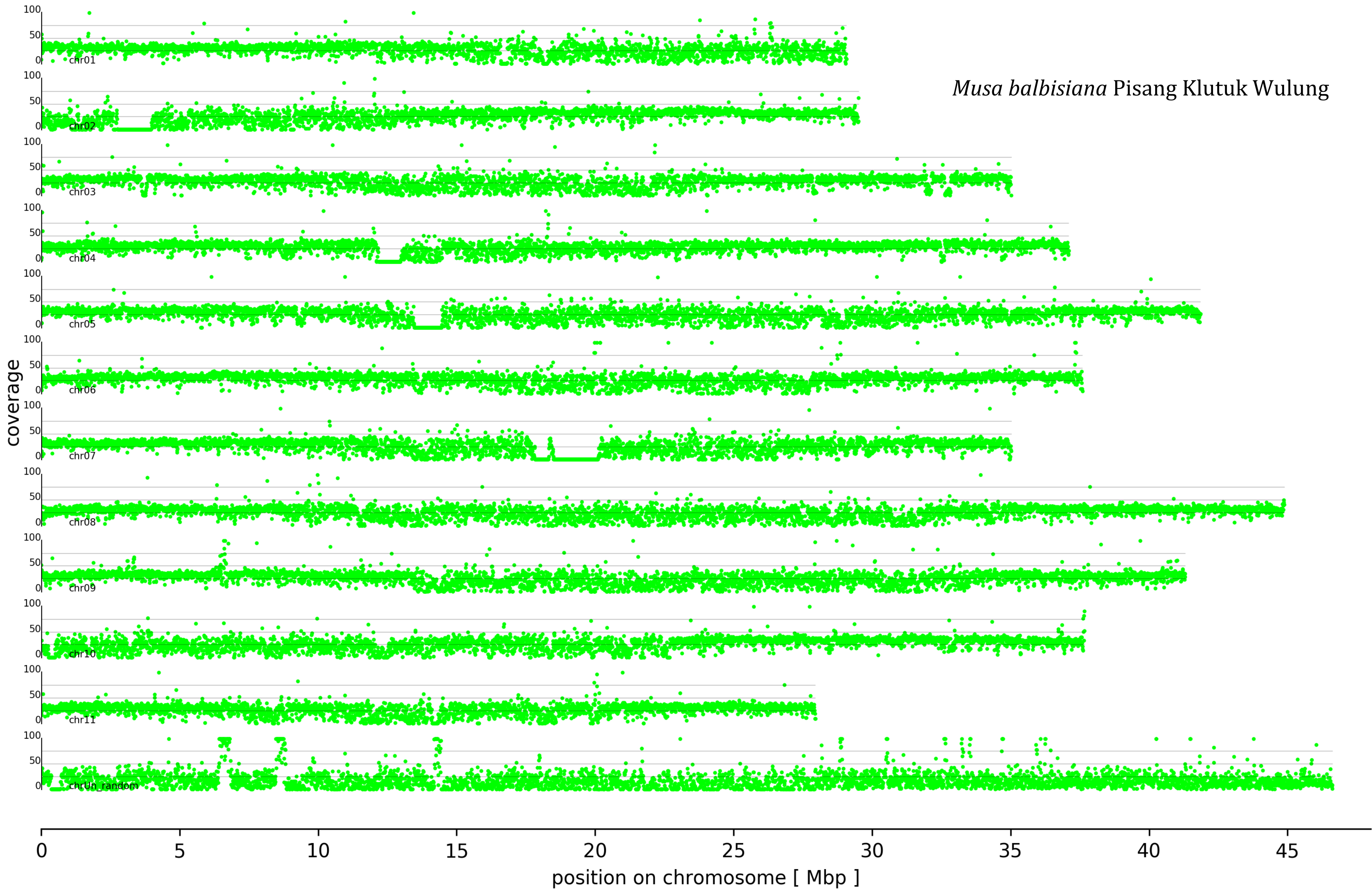
*Musa balbisiana* Pisang Klutuk Wulung
coverage
100
50
0
chr01
chr02
chr03
chr04
chr05
chr06
chr07
chr08
chr09
chr10
chr11
chrUn_random
0
5
10
15
20
25
30
35
40
45
position on chromosome [ Mbp ]

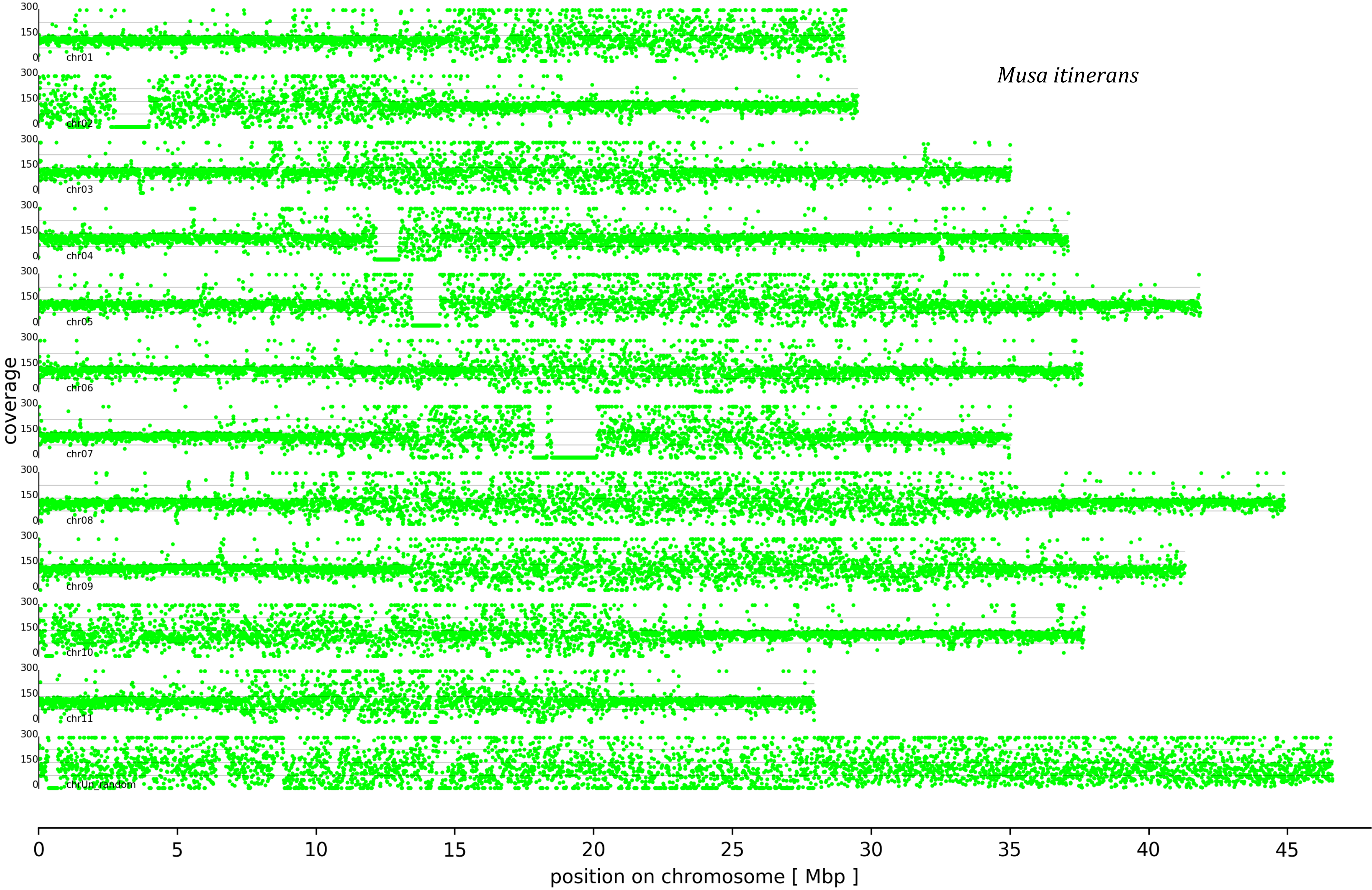

Musa itinerans
coverage
300
150
0
chr01
chr02
chr03
chr04
chr05
chr06
chr07
chr08
chr09
chr10
chr11
chrUn_random
0
5
10
15
20
25
30
35
40
45
position on chromosome [ Mbp ]

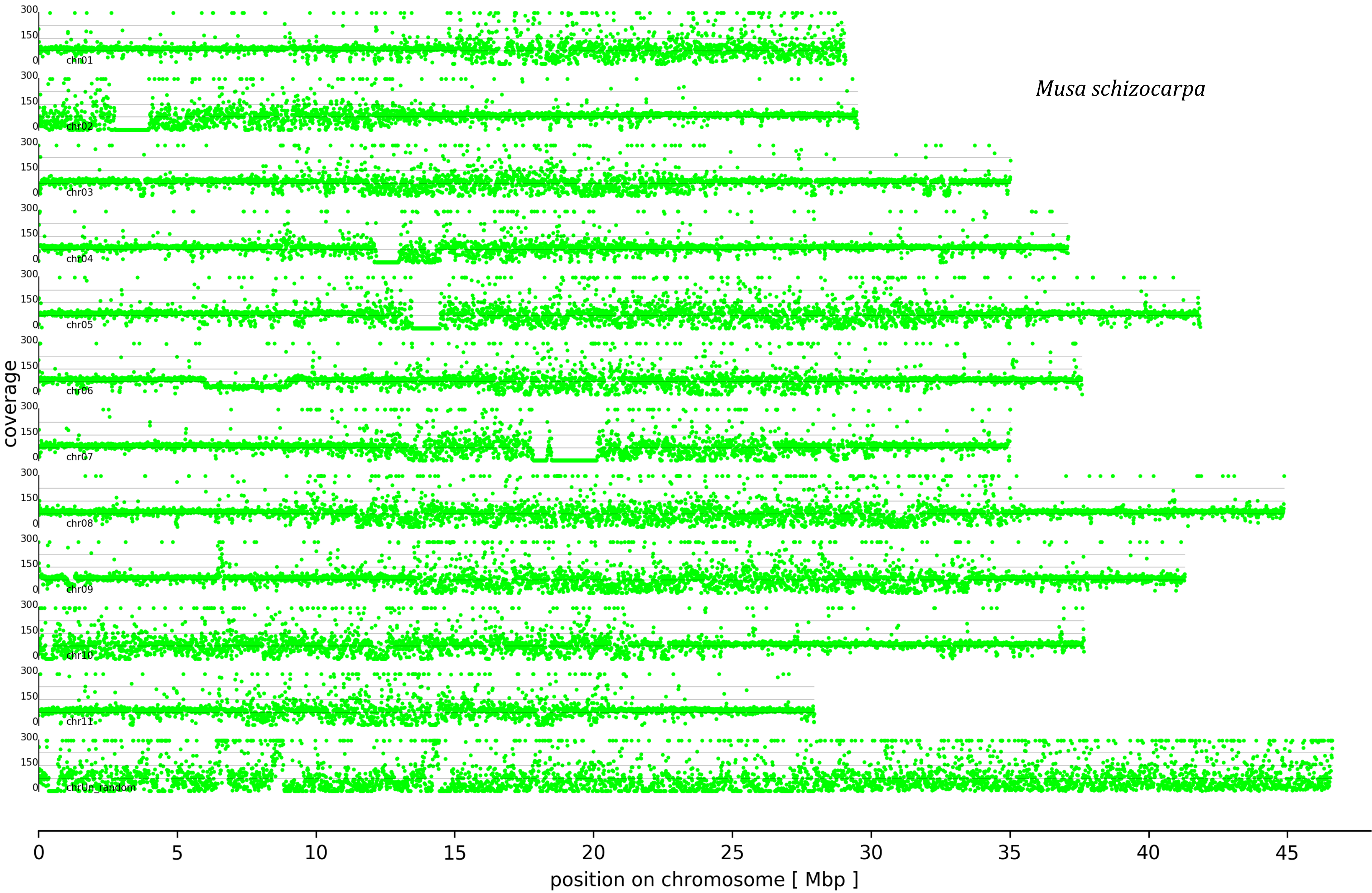
Musa schizocarpa
coverage
300
150
0
chr01
chr02
chr03
chr04
chr05
chr06
chr07
chr08
chr09
chr10
chr11
chrUn_random
0
5
10
15
20
25
30
35
40
45
position on chromosome [ Mbp ]
